# The locus coeruleus is a complex and differentiated neuromodulatory system

**DOI:** 10.1101/109710

**Authors:** Nelson K. Totah, Ricardo M. Neves, Stefano Panzeri, Nikos K. Logothetis, Oxana Eschenko

## Abstract

Understanding the forebrain neuromodulation by the noradrenergic locus coeruleus (LC) is fundamental for cognitive and systems neuroscience. The diffuse projections of individual LC neurons and presumably their synchronous spiking have long been perceived as features of the global nature of noradrenergic neuromodulation. Yet, the commonly referenced “synchrony” underlying global neuromodulation, has never been assessed in a large population, nor has it been related to projection target specificity. Here, we recorded up to 52 single units simultaneously (3164 unit pairs in total) in rat LC and characterized projections by stimulating 15 forebrain sites. Spike count correlations were low and, surprisingly, only 13% of pairwise spike trains had synchronized spontaneous discharge. Notably, even noxious sensory stimulation did not activate the entire population, only evoking synchronized responses in ~16% of units on each trial. We also identified novel infra-slow (0.01-1 Hz) fluctuations of LC unit spiking that were asynchronous across the population. A minority, synchronized possibly by gap junctions, was biased toward restricted (non-global) forebrain projection patterns. Finally, we characterized two types of LC single units differing by waveform shape, propensity for synchronization, and interactions with cortex. These cell types formed finely-structured ensembles. Our findings suggest that the LC conveys a highly complex, differentiated, and potentially target-specific neuromodulatory signal.

---

In contrast to synaptic transmission-based interactions, neuromodulation has long been seen as “one-to-many” activity, with neuromodulatory nuclei often considered to be undifferentiated “state-controllers”. Example *par excellence* is the noradrenergic locus coeruleus (LC), a diffusely projecting brainstem nucleus containing only approximately 1,600 neurons in the rat and 10,000 in humans (per hemisphere). The LC, as a part of the ascending reticular activating system for arousal, is conserved across vertebrates, including teleosts (e.g. zebrafish), amphibians, birds, and mammals, such as rodents and humans ^1,2^ (although neurochemical and neurophysiological differences exist, for example, in cat LC in comparison to the highly similar rodent and monkey LC ^3^). The LC is thought to regulate broad networks related to a multitude of functions, such as autonomic activity, endocrine function, nociception, sleep and arousal, perception, attention, decision-making, learning, and memory ^4^. Its neurons are considered to act synchronously to non-specifically modulate the state of neuronal excitability in many forebrain targets via simultaneous norepinephrine (NE) release ^2,3,5–7^. The seemingly undifferentiated activity of LC has influenced diverse theories ranging from neural control of sleep to computational models of decision making ^5,8–10^.

This perspective of global neuromodulation emerged primarily from two lines of research. The first one comprised anatomical and neurochemical studies demonstrating that the axons of individual LC neurons branch widely to innervate distant forebrain regions where their terminals release NE, which can spread up to ~100 μm in the rodent cortex (volume transmission of NE also occurs in primate cortex although the spread in their larger brain is unknown) ^11–14^. The second line included electrophysiological experiments showing that, in LC, multi-unit activity (MUA) is synchronized with changes in the local field potential (LFP, a marker of transmembrane currents and other peri-synaptic activity within LC), that were registered synchronously with spatially segregated electrodes placed in the core of the nucleus ^2,15,16^. Uniform LC cellular activity is seen as the result of (*i*) shared synaptic input, (*ii*) gap junction coupling, and (*iii*) intrinsic membrane potential oscillations at < 1Hz ^15,17,18^. By firing together, global NE release is thus achieved for the purposes of modulating communication across broad forebrain circuits and for regulating global states of neuronal excitability ^4,7^. In support of this line of thinking, studies of another neuromodulatory system (i.e., dopamine), have also revealed a similarly high degree of population synchrony, consistent with the notion that neuromodulatory neurons broadcast a redundant reward-related, salience, and/or arousal signal ^19–28^.

Although recent anatomical studies have demonstrated that individual LC neurons could provide localized forebrain neuromodulation by targeting different cortical sites ^29,30^, synchronous activity across LC neurons would still result in non-specific neuromodulation. However, prior estimates of synchrony using LC MUA may not be accurate because single units, which could spike independently, have been averaged over. Single unit recordings in LC, though, are rare due to technical challenges. Specifically, the small size of the LC has permitted mainly single wire recordings and the single unit waveforms of the densely packed LC cell bodies are difficult to isolate using a single recording channel. At present, only one study in the awake monkey and one study in the anesthetized rat have managed to simultaneously monitor two single units, and the reported findings were based on small data sets, e.g. ~20 pairs of neighboring single units recorded on the same electrode ^31,32^. According to analysis of cross-correlograms, the spiking of approximately 80% of the unit pairs was synchronized on the timescale of 100 – 200 msec, supporting to the notion of highly synchronized spiking among LC neurons. Evidently, however, recording a small number of pairs (~20) with a single electrode does not allow inferring the degree of synchronicity of a larger LC population. To address this question, we recorded up to 52 single units simultaneously (234 units and 3164 unit pairs in total) using a high-density recording array in urethane-anesthetized rats. In addition, we characterized projection patterns of individual LC units using forebrain electrical stimulation to evoke antidromic responses.

## Identification and characterization of two distinct LC single unit types

We isolated 234 single units in 12 rats and recorded 5 to 52 individual LC units simultaneously (Table 1). Single units exhibited typical electrophysiological and pharmacological characteristics of LC NE-producing cells (Extended Data Figure 1). The extracellular spike waveform shapes of LC single units separated into 2 types based on their spike width and after-hyperpolarization amplitude (Figure 1A, B). We will refer to these populations as “narrow” and “wide” units. Out of 234 single units, 34 units were narrow (15%) and 200 units were wide (85%). Interestingly, beyond distinct spike shape profile, these units had a number of characteristic differences. Narrow units discharged at significantly higher rates compared to wide units (Figure 1C, D; median and s.d.: 1.28±0.73 spikes/sec and 0.64±0.63 spikes/sec, respectively; Wilcoxon-Mann-Whitney, Z=4.23, p = 0.00002, Cohen’s D = 0.823, power = 0.973). The power to detect such an effect size at an alpha level of 0.05 was 97%. Both unit types were distributed throughout the dorsal-ventral extent of the LC, but narrow units were relatively more predominant in the ventral aspect of the nucleus (Figure 1E). This distribution was present in all 12 rats. Both narrow and wide units responded to foot shocks with excitation followed by local NE mediated auto-inhibition, which is typical of LC neurons and all units were inhibited by clonidine, suggesting that both unit types were noradrenergic (Extended Data Figure 1). Moreover, stimulation of forebrain sites elicited antidromic responses in 30% of narrow units and a similar percentage (38%) of wide units (two-sided, Fisher’s Exact Test, Odds Ratio = 1.41, CI = [0.349 5.714], p=0.744), further suggesting that both unit types are likely projection neurons.

**Figure 1.**
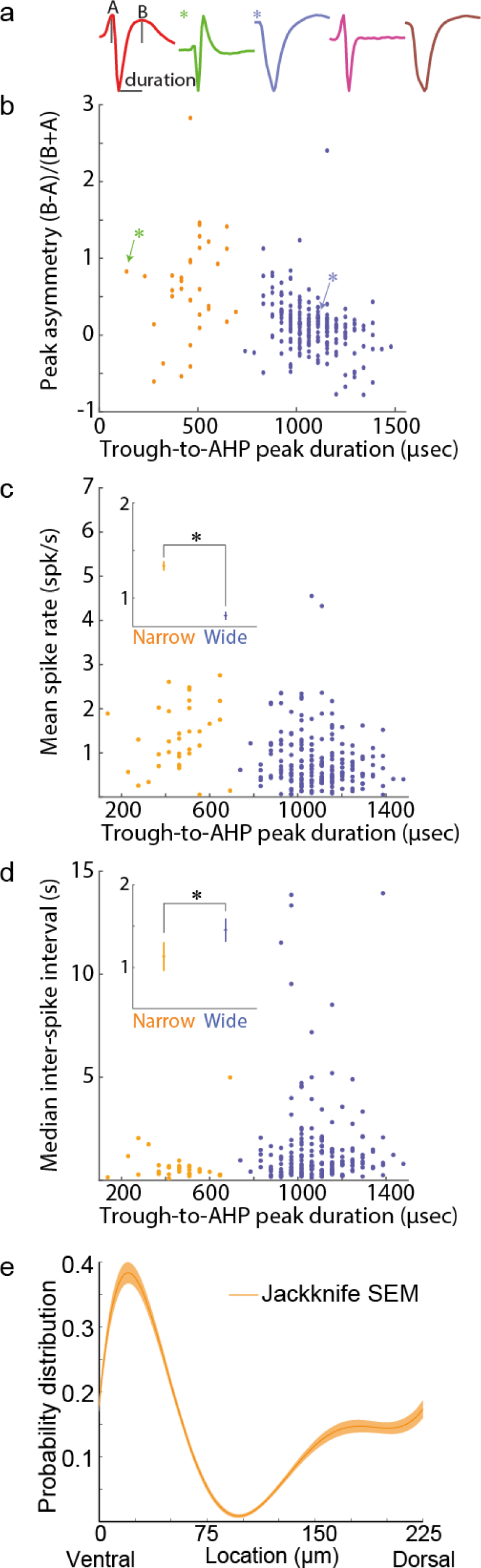
Distinct populations of LC single units were separable by waveform shape, spike rate, and responsiveness to prefrontal cortex stimulation. **(A)** The average waveforms of 5 example units illustrate the diversity of waveform shapes in the LC. **(B)** Units were separable based on the waveform duration and the amplitude of the after-hyperpolarization in relation to the first peak. The green and blue asterisks refer to the example waveforms in panel A with the same markings. **(C, D)** Scatter plots with the mean spontaneous spike rate and inter-spike interval for each isolated LC unit. The insets on C and D show the mean and standard error for each unit type. N = 34 (narrow) and 200 (wide) units. **(E)** Narrow type units were predominantly distributed in the ventral aspect of the nucleus and sparsely recorded elsewhere in the nucleus. The mean and standard error of the probability distribution function is plotted.

**Table 1.** The number of LC single units recorded in each rat. The numbers (and percent) of each unit type are listed. Both unit types were often recorded simultaneously, but narrow units were recorded in 7 out of 12 rats, possibly because of fewer units with narrow waveform reducing their probability of detection

| **Rat Identifier** | **Number of single units** | **Number (%) of narrow type** | **Number (%) of wide type** |
| --- | --- | --- | --- |
| 959.1 | 7 | 5 (71) | 2 (29) |
| 980.4 | 9 | 9 (100) | 0 (0) |
| 105.1 | 5 | 0 (0) | 5 (100) |
| 105.2 | 52 | 10 (19) | 42 (81) |
| 109.1 | 21 | 0 (0) | 21 (100) |
| 134.2 | 15 | 0 (0) | 15 (100) |
| 259.1 | 18 | 0 (0) | 18 (100) |
| 259.2 | 27 | 4 (15) | 23 (85) |
| 263.2 | 36 | 2 (6) | 34 (94) |
| 286.3 | 8 | 3 (38) | 5 (62) |
| 288.1 | 6 | 0 (0) | 6 (100) |
| 288.2 | 30 | 1 (3.3) | 29 (96.7) |

## LC single units have near-zero spontaneous and evoked spike count correlations

While the general consensus is that LC activity is “homogenous”, “uniform” and “synchronous” ^2,3,6,15,16^, the degree of synchrony has not been quantified from a large sample of LC neurons. Systematic large scale recordings in other brain regions have revealed that the degree of synchrony is region-and function-specific ^33^. Knowing where LC falls on this continuum of synchrony would provide insight into the representations and computations of its neurons and the functional role of its output.

Although spike count correlations have been measured for cortical neurons ^34^, they have not been quantified in the LC. Earlier recordings, which were limited to a total of 23 LC neuronal pairs, estimated 80% synchronous discharge by analyzing spike train cross-correlograms and detecting coincidental spiking over 100 – 200 msec epochs^32^. The best estimate of LC synchrony, therefore, greatly exceeds the values obtained for cortical cell-pairs, of which only 16% (32 out of 200 pairs) are reported to have a large central peak in the spike train cross-correlogram ^35^. This yields a correlation coefficients of ~0.05 to 0.1 in cortex ^34^. (Note that, when a spike train cross-correlogram is integrated over sufficiently large lags, the integral approximates the spike count correlation coefficient ^36^). Thus, given the 80% population synchrony that has been reported among LC spike trains, one might expect relatively LC correlation coefficients and the proportion of significant correlations to greatly exceed those in cortex (0.1 and 16%, respectively). In order to establish a quantitative measure of correlated firing in LC, we calculated the spike count correlation coefficient for 3164 LC single unit pairs. We analyzed spike counts in bins of 200 msec and 1 sec, which were each chosen based, respectively, on the 100-200 msec duration of coincidental spiking reported in LC cross-correlograms ^32^ and the previously demonstrated relationship between rhythmic increases of LC multi-unit firing in relation to cortical 1 −2 Hz slow waves ^37,38^. The correlation coefficients were distributed around zero (Figure 2). The mean correlation coefficient across all 3164 pairs of recorded units was 0.044±0.001 for 200 msec bins and 0.098±0.003 for 1 sec bins. Pairwise correlated variability did not depend on a distance between the units (Extended Data Figure 2A). One factor to consider is that higher values of correlations are generally reported under various types of anesthesia and this has been shown to be largely due to the fact that, under anesthesia, neurons are more locked to slow network activity and/or neuronal population oscillations ^39–43^. To ease this concern, we developed a non-parametric permutation test of significance of correlations that discounts the spurious correlations due to non-random inter-spike intervals and common locking to slow oscillations that may arise because of anesthesia. Only 16% of pairs had synchronous spiking that was significantly higher than what would be expected to occur by chance (one-sided permutation test, p<0.01, see Materials and Methods). This suggests that synchrony between LC cells, though present, is much weaker and sparser than expected from previous reports ^32^. Given that correlations tend to be larger when firing rate is higher firing ^44^, one possible concern is that within-LC correlations may be sparser and weaker in anesthetized animals, compared to the awake animal, simply because LC firing rates are slightly lower under anesthesia. To address this concern, we separately analyzed unit pairs with a geometric mean rate that was similar to the rate observed in awake rats and non-human primates (greater than 1 Hz, ^2,45–50^). We found that 19.2% of these higher rate pairs (N = 506) had significantly positive correlation coefficients (one-sided permutation test, p<0.01), which was similar to the percentage (15.8%) of lower rate pairs (N = 2658). Thus, our results show that correlated spiking is not predominant between pairs of LC neurons, even when firing rate is quantitatively similar to those observed in the awake state. The values of correlation coefficients suggest that spontaneous LC population activity may not be described as a purely synchronous pool that broadcasts a homogenous signal.

**Figure 2.**
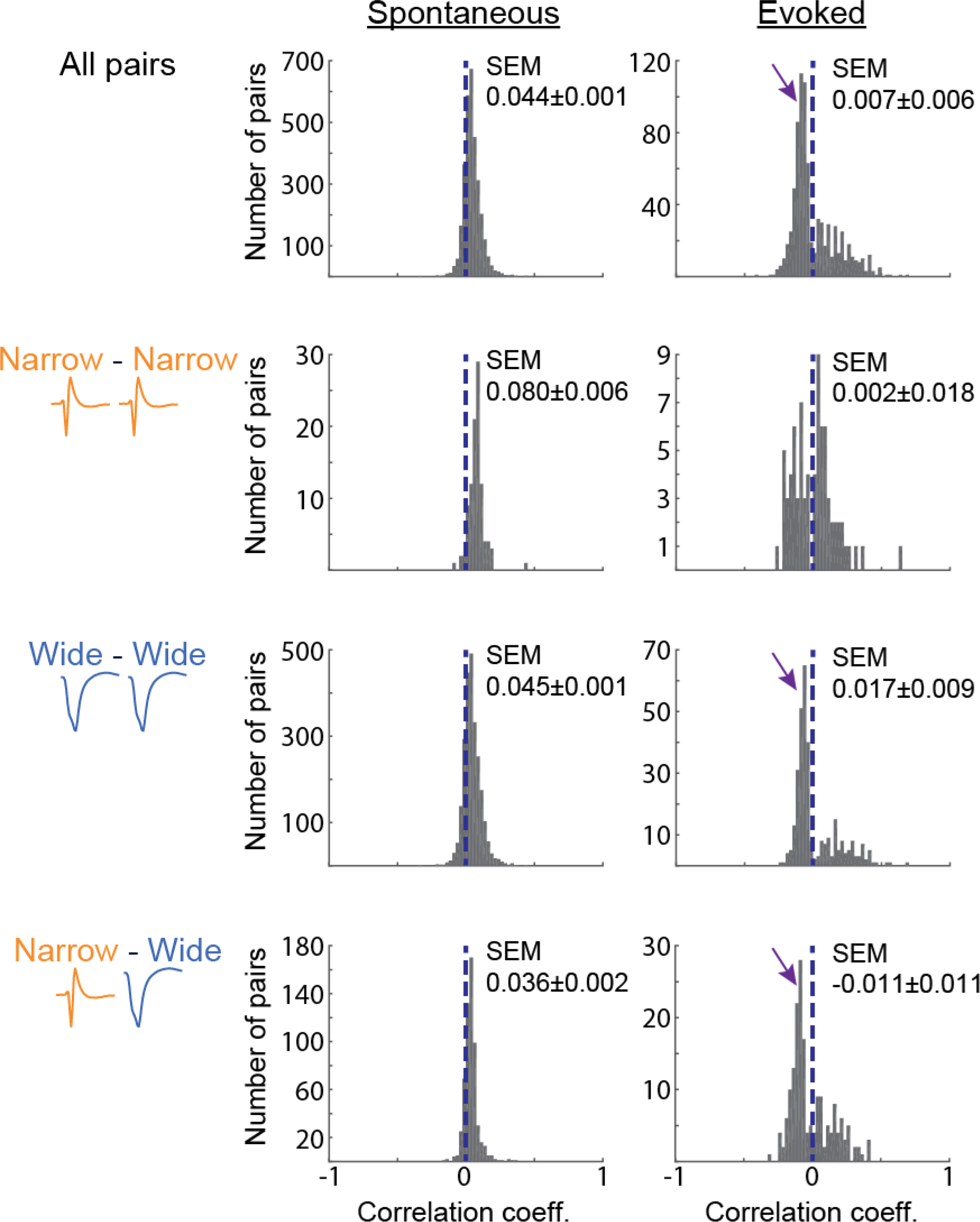
Pairwise spike count correlations are near zero and anti-correlations are cell type specific. The distribution of spike count correlation coefficients is around zero (dotted blue line) during spontaneous spiking (left panel) or following single pulse foot shock stimulation (right panel). Spike count correlations are plotted separately for pairs of narrow units, pairs of wide units, and pairs of mixed unit type according to the labels on the left of the histograms. N = 100 (both narrow type units), 2480 (both wide type units), and 584 (mixed unit type) pairs. Prominent negative correlations are apparent after evoked foot shocks (arrows, right panel). These negative correlations only occur in pairs containing a wide unit.

Sensory stimuli evoke burst spiking of LC neurons, which is thought to be a robust, homogenous population response to each stimulus ^16^. In order to study correlations between evoked LC spiking activity, we applied a single foot shock (5.0 mA, 0.5 msec pulse duration) and measured spike count correlations during the time window of the maximal evoked discharge (50 msec after stimulation). The mean evoked spike count correlation was distributed around zero (Figure 2). Increasing the stimulus intensity to 5 pulses (at 30 Hz) did not increase synchrony (single pulse: 0.007±0.006 versus 30 Hz: 0.012±0.006, Wilcoxon-Mann-Whitney, Z=1.24, p = 0.214). Moreover, accounting for possible adaptation by calculating correlations in blocks of 5 trials (e.g., trials 1 – 5, 6 – 10, and so on) did not demonstrate any propensity for higher correlations during earlier stimulation trials for either stimulation intensity (single pulse: Kruskal-Wallis H = 0.68, p = 0.711; 30 Hz: Kruskal-Wallis H = 0.69, p = 0.709). Thus, the low trial-by-trial evoked spike count correlations suggest that individual units respond independently from each other on every trial. Indeed, within 50 msec after a foot shock a robust population response is easily observed, yet on average only 16% of units responded on each trial of a single foot shock (Figure 3). The proportion of neurons responding to the stimulus in each trial remained relatively low when the post-stimulus window was increased to 100 msec (20% of units) or 200 msec (28% of units). This finding strongly contrasts the prevailing view that many LC neurons respond -in unison -to sensory stimuli in a phasic population burst.

**Figure 3.**
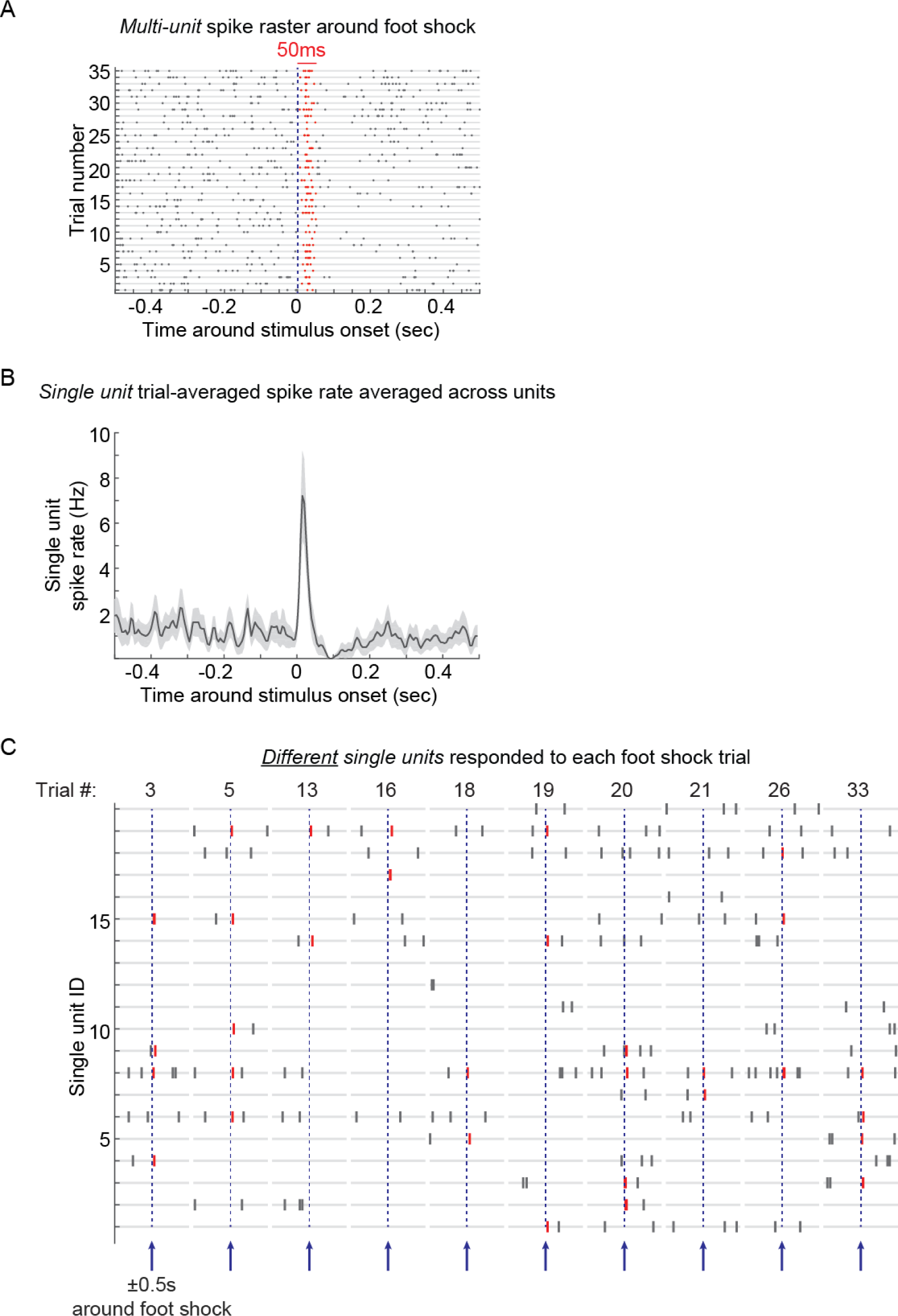
The response to single pulse foot shock engaged different LC units on each trial. The figure shows an example recording of 20 units. **A.** The spike raster (trials are rows) after all simultaneously recorded single unit spike trains were merged into a single multi-unit spike train per rat. The red ticks indicate spiking during the first 50 msec after stimulation (i.e., the population phasic burst of spikes). The phasic response is followed by noradrenergic-mediated auto-inhibition. The plot depicts 20 units from one rat. **B**. The trial-averaged peri-stimulus spike rate histograms for single units were averaged. The plot is the mean and standard error of across single units (N = 20 single units, same as in A, and bin size of 5 msec). This plot shows that single units, on average across trials, exhibit a phasic burst followed by inhibition. **C**. The spike raster of the same 20 units (y-axis) is plotted for 500 msec before and 500 msec after foot shock (arrows) on 10 randomly selected trials. The spikes of each single unit that occur during the 50 msec window that encompasses the LC phasic burst (in A and B) are colored in red. At the level of single trials, different single units responded on different trials. For example, unit 15 responded during the 50 msec after foot shock only on trials 3, 5, and 26; whereas unit 14 responded on trials 13 and 19. This randomness reduces the trial-by-trial evoked spike count correlation coefficients (see Figure 2).

Although the recorded population of LC units exhibited overall weak correlations, we further examined spike count correlations between pairs of narrow or wide units, as well as between unit types. Correlations among each type of pair were similar for spontaneous spiking (Welch’s F(2,253.81)=1.42, p=0.245, ω^2^ = 0.0003). Evoked correlations may differ by unit pair type (but the Kruskal-Wallis test was under-powered, H=7.64, p=0.022, ω^2^ = 0.0003, power = 0.175). Post-hoc tests suggest that pairs of mixed unit type may have a more negative median correlation than pairs of wide type units (p=0.017). Nevertheless, the mean correlation values were near zero for all pair types, which suggests that neither type of LC unit formed a highly correlated sub-population with other units of the same or different type.

Strikingly, a large number of negative spike count correlations emerged during evoked activity. Furthermore, negatively correlated pairs were observed only when the pair included a wide unit (Figure 2, **arrows**). Negative spike count correlations, specifically after sensory stimuli, may reflect lateral inhibition, which is generated by somatic release of NE that inhibits neighboring neurons via alpha-2 adrenoreceptors ^51–53^. Somatic release requires the high frequency of spiking typically associated with sensory stimuli and not spontaneous activity ^54^. Given that both unit types were responsive to salient stimuli, our results suggest that only the stimulus-evoked discharge of wide units generates sufficient somatic NE release to cause local lateral inhibition, but both unit types are noradrenergic and susceptible to lateral inhibition.

## Synchrony due to putative gap junctions or common synaptic input is rare

The presence of a minority of pairs with highly positive spike count correlations suggests that at least some LC single units are correlated. Their synchronized activity could be due to synaptic drive shared by a neuronal pair or electrotonic coupling between the pair ^55–59^, which are both prevalent forms of connectivity in the LC. We assessed the duration of coincidental spiking between unit pairs by measuring cross-correlograms between spike trains. All cross-correlogram analyses used spontaneous spiking. We chose to study coincident spiking on two timescales, tens of milliseconds ("broad-type" interactions) or sub-millisecond ("sharp-type" interaction) that could reflect either common synaptic input or gap junctions, respectively. Shared synaptic input from a third neuron (or group of neurons) appears, instead, as a zero-centered peak which is broad (spread over tens of milliseconds) ^57^. Gap junctions are associated with a sharp peak that is shifted 0.5 to 1 millisecond from zero ^58–60^. We observed coincident spiking on both timescales (Figure 4A). Cross-correlograms were assessed against a spike time-jittered surrogate (grey and blue lines, Figure 4A).

**Figure 4.**
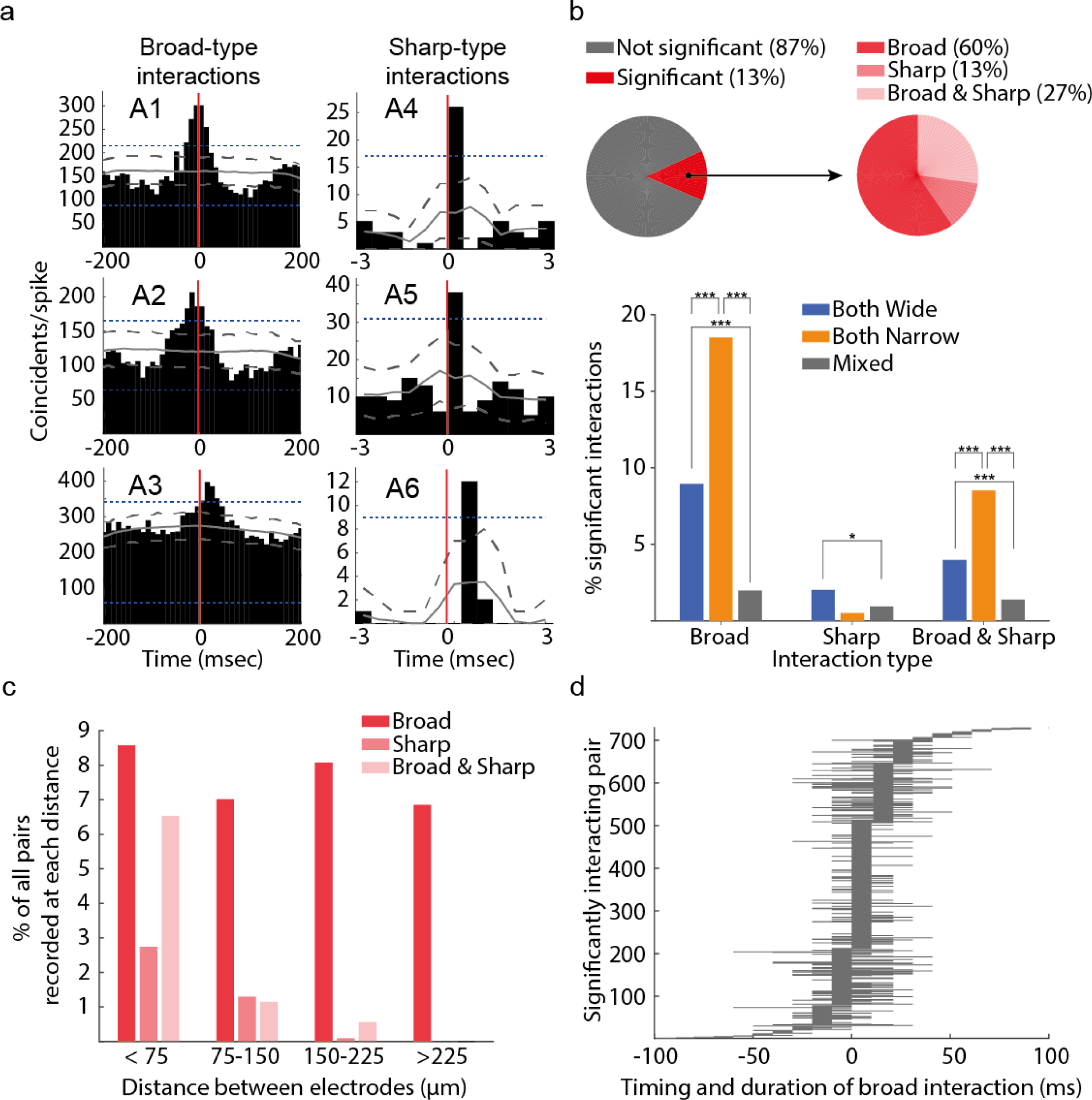
Spike train cross-correlograms indicated that interactions were rare. (A) Three example cross-correlograms with significant coincidental spiking on the timescale of broad-type interactions (A1-A3) and sharp-type interactions (A4-A6). Broad-type interactions lasted for tens of milliseconds, whereas sharp-type interactions were extremely brief (less than 1 millisecond). An interaction was counted if a significant number of coincidental spikes crossed both a pairwise 1% threshold (dotted grey lines) and a 1% global threshold (dotted blue lines) obtained from a surrogate data set of 1000 cross-correlograms computed from jittered spike times. The mean of the 1000 surrogate cross-correlograms is a solid grey line. **(B)** A minority of cross-correlograms (13%) had a significant number of coincidental spikes in at least one bin. A larger percent of the pairs of narrow units had significant broad-type interactions in comparison to pairs of wide units or pairs of both unit types. **(C)** The percent of pairs with broad-type interactions was similar regardless of the distance between the units in the pair. On the other hand, sharp-type interactions occurred only between spatially confined units. **(D)** Out of the pairs with significant broad-type interactions, the timing of the interaction peak and the duration (continuous bins above the significance threshold) is plotted for each pair. These interactions occurred over a broad time range. Peaks were not exclusively centered at time 0 with a symmetrically spread around time 0 (as in panel A1).

We found that only 13% of unit pairs had significant cross-correlations (Figure 4B). Of those 13% of correlated unit pairs, 60% had broad-type interactions, while 13% had sharp-type interactions, and the remaining 27% had both broad-and sharp-type interactions. Thus, only 11% of all 6,299 recorded pairs (interactions considered in both directions) spiked within tens of milliseconds, which is remarkably low, given an estimate of 80% of LC pairs spiking synchronously at this timescale based on prior evidence from 23 pairs ^31,32^. The proportion of synchronized LC neurons is similar to the proportion of synchronized cortical neurons reported as 3.6%, 13%, and 56% under various conditions ^55,61,62^.

A significantly larger proportion of narrow unit pairs had significant broad-type interactions in comparison to pairs of wide units and pairs of mixed unit types (Figure 4B). This finding is consistent, in general, with more positive spike count correlations between narrow units (Figure 2). Furthermore, pairs with broad-type interactions had higher spike count correlations in comparison to pairs with sharp-type interactions and pairs without significant cross-correlations (Extended Data Figure 3). These results are consistent with broad-type interactions and spike count correlations both reflecting common synaptic input. Broad-type interactions, just like spike count correlations, did not depend on the distance between the unit pairs and therefore occurred with similar frequency throughout the LC nucleus (Figure 4C). The distance-invariance of correlated activity in LC (Extended Data Figure 2, Figure 4C) concurs with anatomical evidence of many LC neurons integrating broad and non-topographically organized afferent inputs to the nucleus ^63^. Our findings of little correlated activity suggest the potentially synchronizing influence of shared synaptic input from a broad set of afferents is somehow limited.

We next assessed the dynamics of broad-type interactions by examining the peak times of the cross-correlograms for pairs with significant interactions. In the example cross-correlograms (Figure 4A), there is a notable diversity in the timing of the interaction in different pairs. The interaction in **Figure 4A1** was centered at 0 msec, while interactions between other pairs occurred before 0 msec (**Figure 4A2**) or after 0 msec (**Figure 4A3**). Across the population of all pairs with significant broad-type interactions, the cross-correlogram peak times were spread over ±70 msec (Figure 4D). The peak should be centered at 0 msec if common synaptic input jointly drives the pair ^57^. Therefore, neuronal interactions in LC at this timescale may reflect common synaptic input interacting with other mechanisms that introduce a delay. For example, lateral inhibition ^51,53^ between units sharing a synaptic input could delay their correlated synaptic responses. The local NE release due to discharge of single LC neuron (or a small number of neurons) inhibits spiking of neighboring LC neurons for ~100 msec ^53^. The duration of lateral inhibition demonstrated in this prior work corresponds with the delays observed here for broad-type interactions (Figure 4D), suggesting that lateral inhibition may be responsible for correlated spiking with a delay.

In addition to brief (sub-millisecond to tens of milliseconds) interactions, synchrony could conceivably occur over multiple seconds or even minutes to hours, given that LC spiking is related to arousal ^2^. We used a data-driven approach to potentially detect synchrony occurring in long windows ranging from 20 milliseconds to 40 seconds ^36^. Based on this analysis, we examined cross-correlograms over a ±20 sec window (against a surrogate of jittered spikes). However, out of the already limited set of synchronous pairs observed, the vast majority of synchronous spiking occurred in a window of 70 msec (Extended Data Figure 4). Thus, the time windows we have explored throughout these analyses are sensitive to the timescale of synchrony in the LC.

## Sharp-type interactions are spatially localized

Out of the 13% of cross-correlograms that were significant, 40% were sharp-type interactions, which is 5% of all recorded pairs. These sub-millisecond interactions between LC units fell off rapidly with the distance between units, which may suggest a dependence on electrotonic coupling (Figure 4C). A predominant view of LC function is that gap junctions spread synchrony throughout the LC using collections of electrotonically-coupled neurons ^5,17,32^. Considering the possibility that sharp-type interactions may reflect gap junction coupling (as others do for cross-correlograms of spike trains recorded in the retina, cortex, and cerebellum, ^58–60^), we counted the number of units which exhibited sharp-type interactions with one or more other units. Out of the units with at least one sharp-type interaction with another unit, 38% interacted with only this one other unit (i.e., a network of 2 units), 35% interacted with 2 other units, 16% interacted with 3 other units, and the remaining 11% interacted with 4 to 6 other units. These findings suggest that synchrony on the timescale of putative gap junctions is primarily limited to networks of 2 to 3 units, but also as many as 7 units.

We assessed the propensity of these networks to produce repeating patterns of spiking over a few milliseconds, which could be mediated by putative gap junctions (such that some of unit A’s spikes would be consistently followed by unit B spiking around 1 msec later, followed by unit C spiking around 1 msec after that). Repeating patterns occurred with negligible frequency. In 2 out of 12 rats, we observed triplets of units that spiked in a consistent order over 4 msec (allowing for 0.4 msec jitter of each spike). Only 1 triplet out of 22,100 possible triplet patterns (0.005%) was found in one rat and 4 triplets out of 1,330 possible triplet patterns (0.301%) were found in the other rat. Patterns beyond triplets were never observed. Sharp-type interactions (possibly reflecting electrotonic coupling) are, therefore, spatially limited.

## Spiking of individual LC units oscillates asynchronously at low (< 2 Hz) frequencies

Synchronized rhythmic spiking of LC units could emerge from entrainment with cortical oscillations. In the cortex, these oscillations are prominent during slow wave sleep and anesthesia (but also during the awake state) and include a 1 - 2 Hz "delta oscillation" regime and <1 Hz "infra-slow oscillation" regime ^64–70^. LC MUA has been shown to oscillate in these frequency bands and phase lock to the cortical oscillation leading to the impression that the majority of LC neurons spike together, entrained with the cortical oscillation ^37,38,71^.

We first characterized oscillations in the firing rate of LC single units by calculating the power spectrum of each unit’s spike train converted into a continuous spike density function (SDF, convolution with a 250 msec Gaussian kernel). We calculated the power spectrum of each single unit SDF and then examined the average power spectrum across all 234 single units (Figure 5A). We observed peaks in the infra-slow frequency band. These peaks reflect rhythmic fluctuations in spike rate that are predominant in many single units, but not necessarily synchronous across units. Surprisingly, in contrast to the infra-slow band, we did not observe any distinct peak in the delta oscillation frequency band. This result is unexpected in light of previous studies, which have found that LC multi-unit spike rate oscillated in this range ^37,38,71^. In order to understand the relationship between the activity of LC single units and cortical delta oscillations, we first compared our results with prior studies of LC multi-unit spike rate by merging all simultaneously recorded spike trains into a single multi-unit spike train and converting that to a SDF (250 msec Gaussian kernel). In line with previous studies, we observed that LC multi-unit spike rate did oscillate in the delta frequency band (Figure 5B). Out of 8 rats with spiking during cortical delta oscillations, all 8 of the multi-unit signals were significantly phase locked (Rayleigh's Z test, p<0.05) to the cortical LFP delta oscillations (Figure 5C). Our results demonstrate that if only multi-unit spiking is measured (as is typical in LC recordings), the data suggest that LC neurons respond synchronously along with cortical oscillations; however, our data reveal that this is not the case at the single unit level. In spite of LC single units not exhibiting spike rate fluctuations at ~ 1 – 2 Hz (Figure 5A), approximately 69% of single units were significantly phase locked to the cortical delta (Rayleigh’s Test for Circular Uniformity, p<0.05). Individual LC neurons, therefore, respond during periodic (1 - 2 Hz) transitions to states of heightened cortical excitability, but each on different cycles rather than as a synchronized, rhythmically fluctuating population which yields no 1 - 2 Hz peak in the single unit power spectrum.

**Figure 5.**
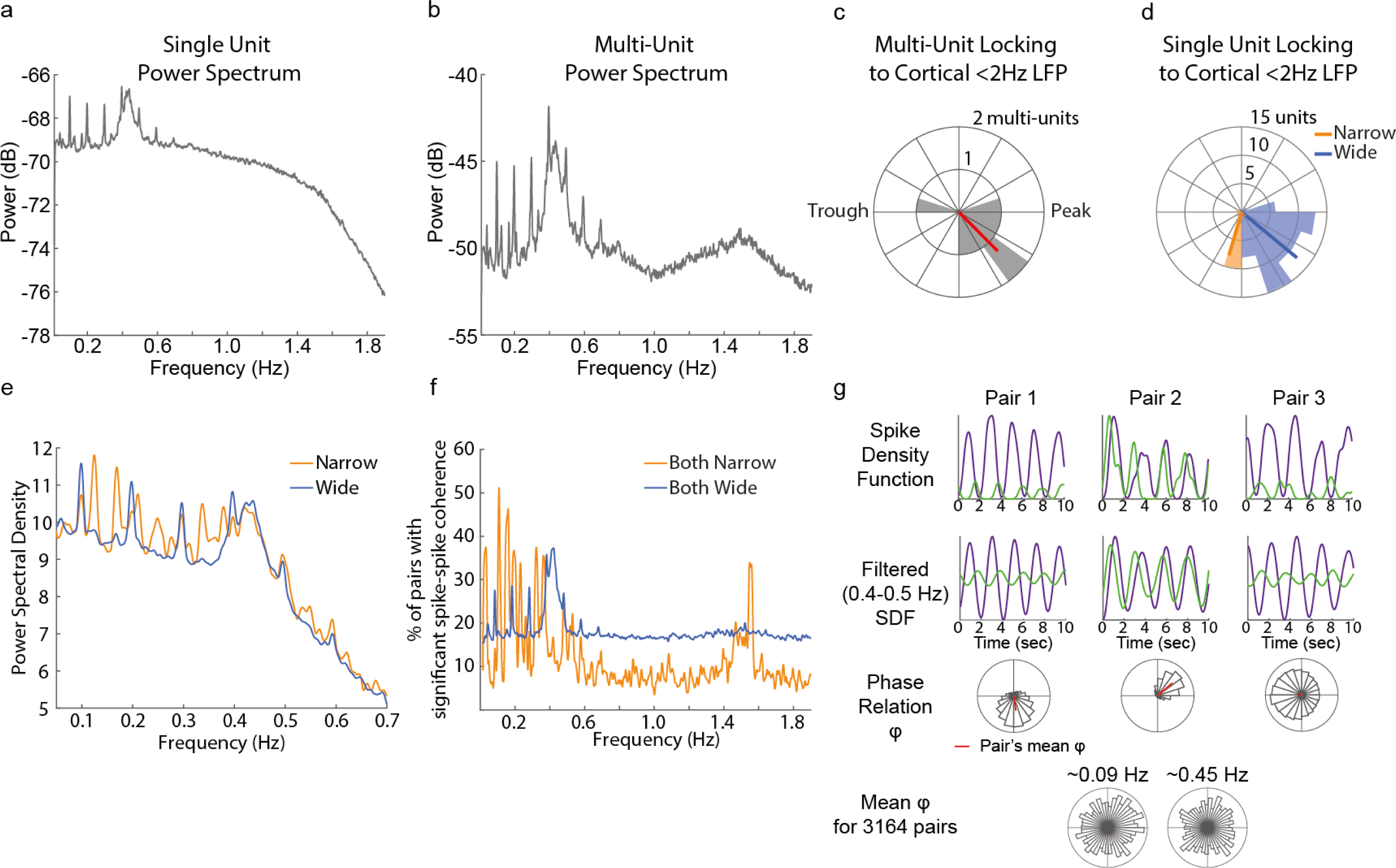
Spike rates oscillated asynchronously across individual LC units and different cell types responded at distinct phases of cortical slow oscillations. **(A)** Single unit spike trains were converted to spike density functions (SDF) and their power spectra were calculated. The plot shows the mean power spectrum across all single units. Two oscillatory regimes at 0.09 Hz and between 0.4 and 0.5 Hz were observed in single unit spike trains. **(B)** Merging simultaneously recorded single unit spike trains into one multi-unit spike train before constructing a SDF revealed spectral power at the infra-slow frequencies (0.09 Hz and 0.4-0.5 Hz), as well as at around 1 – 2 Hz. **(C)** Multi-unit spiking was phase locked to the trough-to-peak transition reflected by the phase of cortical delta oscillations. The polar plot is a histogram of the number of multi-units at each bin of cortical LFP phase. The red line is the mean across multi-units. **(D)** Single units were also phase locked to the cortical delta oscillation, in spite of not rhythmically spiking at that frequency (A). Narrow units responded significantly earlier than wide units during the cortical delta oscillation. The wide units preferentially fired at 320 degrees, whereas the mean angle for narrow units was 254 degrees (the trough, or down state, was 180 degrees and the zero-crossing between the trough and the peak was at 270 deg). **(E)** Wide and narrow units both oscillated at infra-slow frequencies (0.09 Hz and 0.4-0.5 Hz), but narrow unit spike trains had additional peaks of spectral power between 0.1 and 0.2 Hz. **(F)** The spike counts of narrow and wide units fluctuated coherently at a range of infra-slow and slow frequencies. The percent of pairs with significant spike-spike coherence is plotted for pairs of narrow units and pairs of wide units. Pairs of wide units oscillated together at 0.09 Hz and 0.4-0.5 Hz. Pairs of narrow units oscillated together at different infra-slow frequencies and at approximately 1.5 Hz. **(G)** The spike rates of 3 example pairs over a 10 sec period are plotted as spike density functions (top panel) and filtered for 0.4-0.5 Hz (middle panel). The bottom panel is a histogram of the phase differences (ϕ) between the units’ oscillations over the entire recording session. The units in each pair oscillated coherently with narrow distributions of phase differences, but only the units in Pair 2 oscillated nearly synchronously (i.e., with ~ 0 degrees of phase difference). The mean ϕfor each example pair is marked by the red line on the polar plot. Note that the phase angle histograms and mean ϕillustrate the stable phase consistent relationships between pairs over the entire recording (often multiple hours), not only the 10 sec plotted in the above panels. At the population level of all 3164 pairs, the mean ϕacross pairs are distributed uniformly across all phases. A pair was included in the population histogram if its distribution of phase relations was significantly non-uniform (Rayleigh’s Test for Circular Uniformity, p<0.05), as in the 3 example pairs.

Similar proportions of each unit type (70% of wide units and 66% of narrow units) were phase locked (two-sided Fisher’s Exact Test, Odds Ratio=1.23, CI=[0.470 3.202], p=0.803). Notably, narrow units responded significantly earlier in the cortical delta oscillation (Watson-Williams test for equal circular means, F(83)=40.959, p<0.0001; Figure 5D). Narrow units responded during the LFP delta oscillation trough to peak transition, while wide units responded closer to the LFP peak; thus, in contrast to the canonical thinking that LC neurons act homogenously to precipitate up states, we show that each unit type may make differing contributions to neuromodulation of cortical excitability ^7,37,38^.

Intriguingly, we observed strong single unit spike rate oscillations in the infra-slow frequency band (Figure 5A), specifically at 0.09 Hz (periods of 11 sec) and 0.4 – 0.5 Hz (periods of around 2 sec), which were readily observable in SDF's (Figure 5G, **top panel**). Additional examples for the 0.09 Hz oscillations are presented in Extended Data Figure 5. Both unit types oscillated at these frequencies and narrow units also oscillated at additional frequencies between 0.1 and 0.2 Hz (Figure 5E). The infra-slow oscillations were coherent between pairs of units (Figure 5F). Strong coherence between unit pairs suggests that synchronous spiking of LC unit pairs may occur at infra-slow oscillatory time scales. Therefore, we next examined the phase relationship of the spike rate oscillations between LC units in the infra-slow range. Spiking of the majority of pairs (73% for 0.4 - 0.5 Hz and 67% for 0.09 Hz) oscillated coherently with a stable phase relationship (Rayleigh's Test for Circular Uniformity, p<0.05). The three examples exhibited stable phase relationships, with one pair's spiking fluctuating synchronously (in-phase at nearly 0 degrees phase difference), whereas other pairs responded in a stable anti-phase pattern (180 degrees phase difference) such that their spiking was consistently in opposition (asynchronous) over infra-slow time scales (Figure 5G, bottom panel). At the population level (all 3,164 pairs), the mean phase relations across all pairs were distributed uniformly for spike rate oscillations at both 0.09 Hz (Rayleigh’s Z=2.531, p=0.080) and 0.4 – 0.5 Hz (Rayleigh’s Z=1.074, p=0.342). These data indicate that most pairs exhibit coherent oscillations, but only a small portion oscillate synchronously (in-phase), yielding little oscillatory synchrony at the whole population level.

## LC single units exhibit complex population patterns and form ensembles

Although we have found multiple lines of evidence that LC single units do not respond synchronously, it is in principle possible that the LC contains smaller groups of units with synchronized activity, that is, cell ensembles. For example, we observed a minority of highly correlated unit pairs (long right tails in the spike count correlation coefficient distributions in Figure 2 and 13% of pairwise cross-correlograms were significant in Figure 4). To explore this further, we measured the coupling of single unit spiking to the spiking of the population (all remaining units) with 1 msec resolution. Population coupling measures the number of spikes that occur in the population in the same msec as a single unit spike ^72^. During spontaneous activity, population coupling varied across individual single units. For example, the spiking of example Unit A was highly synchronous with other units in the population (Z-score at time 0 is ~13), whereas example Unit B was uncoupled (Z < 2 at time 0) from the population (Figure 6A). The distribution of Z-scores at 0 msec across all single units indicated the presence of both uncoupled single units (34% of units had Z<2) and population coupled units (Figure 6B). Population coupling suggests that some sub-sets of multiple units may be synchronously active as ensembles. The one millisecond timescale of population coupling suggests that ensembles may be active on extremely brief scales.

**Figure 6.**
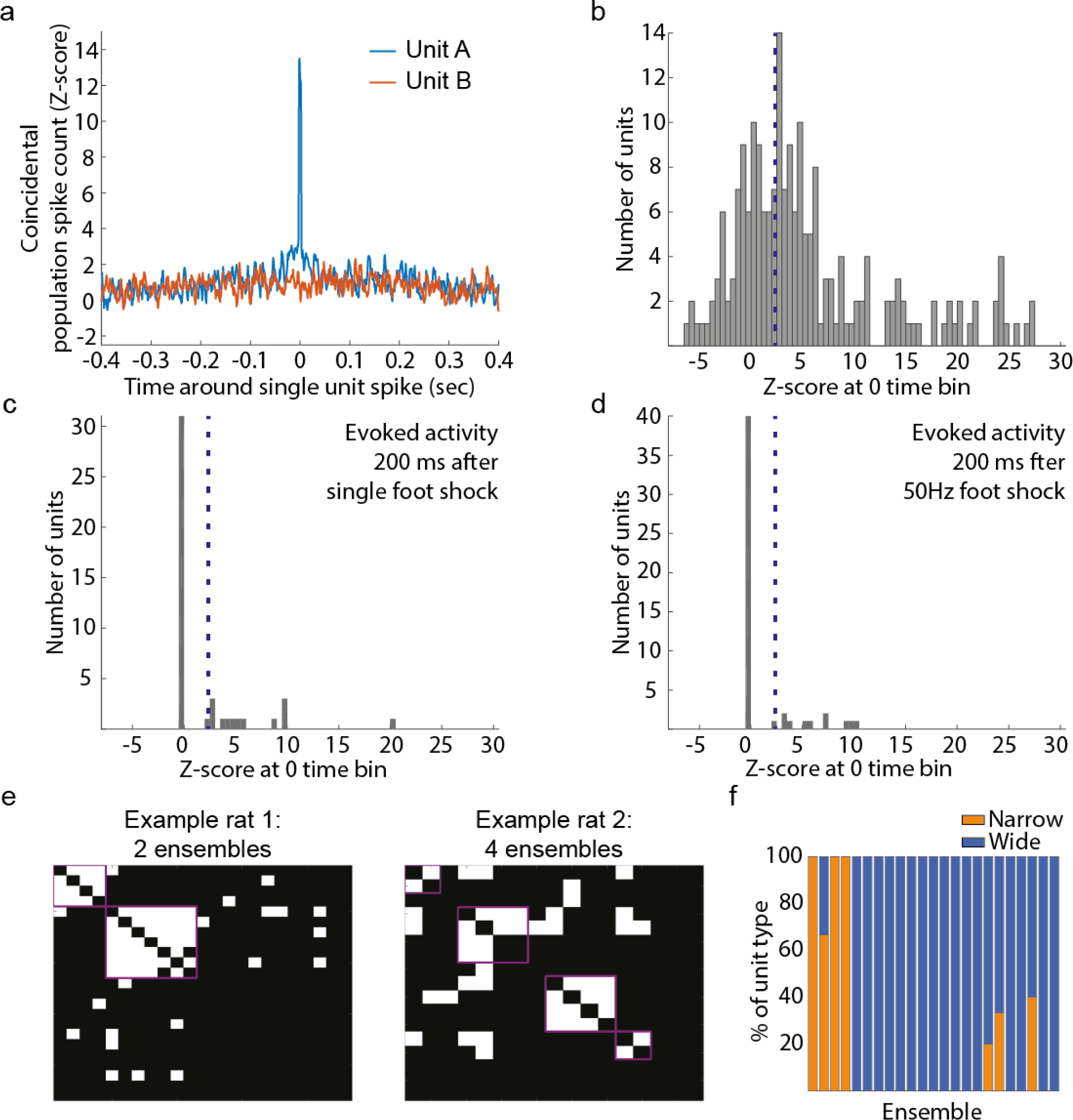
LC single units have diverse coupling to the population and organize into
ensembles of homogenous unit type. **(A)** Population coupling was calculated as the cross-correlogram between each single unit and the merged spike train of all remaining single units that were simultaneously recorded. The cross-correlogram was calculated with 1 msec bins over a period of ±400 msec and Z-score normalized to the period between 300 and 400 msec (edges of the cross-correlogram). Example unit A illustrates a case of strong population coupling. Spikes of unit A coincided with spikes of many other units recorded simultaneously. Example unit B illustrates a lack of population coupling. **(B)** A histogram of the population coupling strength (Z-score at time 0) illustrates the broad range of population coupling strength across LC single units. Dotted lines show a Z-score of 2 for reference. **(C, D)** Population coupling using spikes during the 200 msec after a single foot shock (C) or after a brief train of foot shocks (5 mA pulses at 30 Hz) (D). As with spontaneous activity, many single units are not coupled to the population, with the exception of those units on the right tail of the distribution. **(E)** Two examples of ensembles detected in two rats. White dots indicate correlated units. Magenta lines outline ensembles. Ensembles are defined as correlated activity between two or more single units. All simultaneously recorded single units were treated as a network with links between correlated pairs and ensembles were detected using community detection algorithms on the network. **(F)** The percent of each unit type making up each ensemble. Each bar is one of 23 ensembles. The majority of units in each ensemble were the same type.

Sensory stimulation is thought to evoke synchronous discharge of many LC neurons ^16^ and should therefore result in strong population coupling for most single units. Astonishingly, foot shocks did not cause coupling of a large number of single units to the population (Figure 6C, D), suggesting a lack of synchronous population discharge to sensory stimuli at a msec time scale, in line with our earlier pairwise analysis (Figure 2B, 2C, Figure 3).

We next attempted to detect and discriminate which LC units formed correlated sub-populations spiking together as ensembles using graph theory analysis. We observed ensembles in each set of simultaneously recorded units (Figure 6E). We identified a total of 23 ensembles, ranging from 1 to 3 per rat, and consisting of 2 to 9 units per ensemble. Ensembles were most likely due to distance-invariant shared synaptic inputs (Extended Data Figure 2, Figure 4C), which contributed the majority of correlated activity in the nucleus; correspondingly, LC unit ensembles were spatially diffuse (Extended Data Figure 6). Surprisingly, the units in an ensemble often consisted of the same unit type (Figure 6F).

## A minority of correlated single units provide targeted forebrain neuromodulation

We examined the degree to which correlated units have overlapping projection targets. We assessed the projection properties of LC cells by applying direct electrical stimulation in up to 15 forebrain sites. Example spike rasters showing antidromic responses, latencies to respond, number of projection targets, and firing rates of units based on projection target are shown in (Extended Data Figure 7). The mean latencies for each projection target are consistent with the prior literature ^73–75^. In general, positively or negatively correlated unit pairs (those with significant correlation at p<0.01, permutation test) did not have any greater tendency for both units to jointly project to overlapping forebrain targets (Figure 7A, B). Additionally, pairs with broad-type interactions (assessed by spike train cross-correlograms) did not relate to the degree of target overlap between units (Figure 7C). However, pairs with sharp-type interactions were more likely to project to the same target (Figure 7D). Significantly more pairs with sharp-type interactions (88%) than non-interacting cell pairs (72%) had overlapping projections to the same forebrain zone (Cohen’s D = 0.55, one-sided Fisher’s Exact Test, p=0.075). Out of the unit pairs with overlapping projection zones (e.g., any division of prefrontal cortex or sensory cortex or thalamus), cell pairs had a similar tendency to project to cortex more than thalamus and prefrontal cortex over sensory cortex (Figure 7D, **inset**).

**Figure 7.**
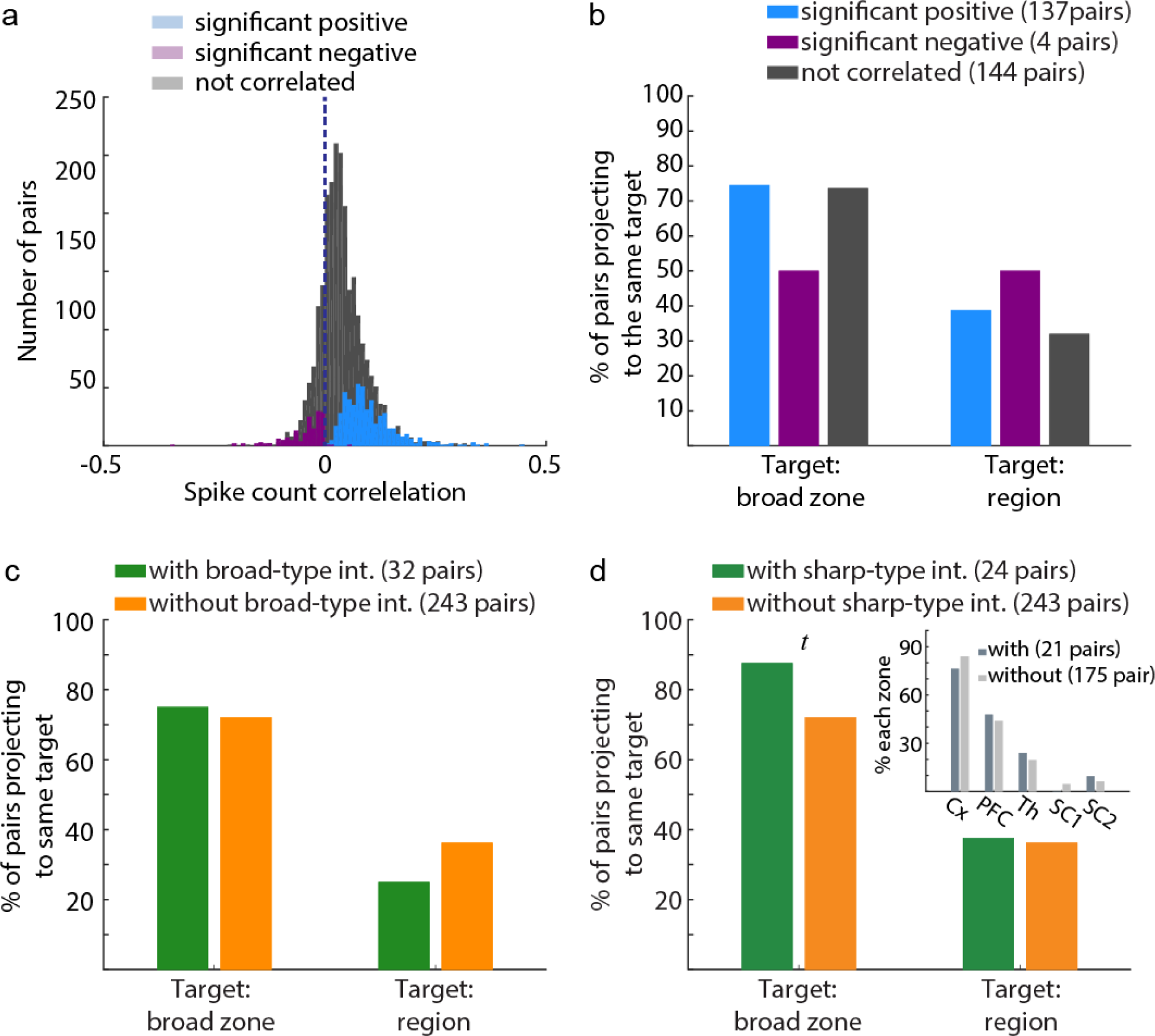
The minority of unit pairs with synchrony on the timescale of gap junctions provided targeted forebrain neuromodulation. **(A)** Spike count correlations coefficients were divided into highly correlated pairs (Pearson’s correlation coefficient, p<0.001) with positive (blue) or negative (purple) correlations. **(B)** The percent of pairs with both units jointly projecting to the same forebrain target did not differ between pairs with correlated activity (blue), anti-correlated activity (purple), and non-correlated units (grey). The percent is out of the total number of units indicated in the figure legend. Targets were defined as either individual brain regions or as zones (i.e., cortical, sub-cortical, thalamic, prefrontal, primary sensory cortex, or secondary sensory cortex). Zones were examined because prior work has indicated that single LC neurons may project to multiple, functionally-related forebrain sites ^114^. **(C)** The percent of pairs with overlapping projection targets did not depend on the pair having a network interaction. **(D)** Pairs with gap junction interactions had a significantly greater likelihood of projecting to the same forebrain target zone.

## Discussion

### The LC contains multiple, functionally differentiated cell types

We observed two types of LC units that differed by waveform shape, firing rate, propensity for synchronization, and interactions with cortex. The cell types also had remarkably different dorsal-ventral distributions within the LC. Our findings provide at least three lines of evidence that these cell populations may have different functions. First, we found that each cell type has unique local circuit properties, namely narrow and wide units have differing capabilities to cause local NE-mediated lateral inhibition and oscillatory firing rate changes (spike-spike coherence) at different frequencies. Second, we revealed that each cell type tended to form ensembles with other cells of the same type. Third, we observed differential phase locking to cortical oscillations, which may imply distinct roles in modulating cortical excitability. This latter finding suggests that the activity of LC narrow units (which were locked to an earlier phase of cortical LFP delta oscillations and had a higher firing rate that should release more NE) could function as a cell type-specific mechanism for regulating cortical excitability in a more selective manner than is commonly attributed to neuromodulatory systems ^7^. Although anesthesia also alters cortical LFP ^76,77^, delta oscillations appear very similar in the awake, sleeping, and anesthetized rodent ^64–66^. Further investigation of the relationship between cortical LFP and LC cell types during natural states of wakefulness and sleep is warranted now that these LC cell types have been described.

The significance of multiple LC cell types remains to be seen. Although their morphological, genetic, and membrane electrophysiology characteristics cannot be resolved with extracellular recordings, our work reveals a new level of diversity among LC single units. This diversity introduces many outstanding questions -- from the local circuit and afferent connectivity principles that underlie formation of LC cell type-specific ensembles -- to how LC ensemble activity patterns modulate forebrain targets. Recent work suggests that separate LC ensembles may modulate distinct forebrain targets to control different behaviors ^78^.

### Population synchrony occurs among a small proportion of LC neurons

Correlations have been consistently reported between neuronal pairs within many brain structures and are thought to be a defining feature of neural population activity. The strength of correlation varies across brain regions and may have profound influences on the time scales of the computations that neuronal ensembles perform and the functional relevance of their output ^33^. The most prominent hypothesis is that the LC ^2,3,5,6,22^ and, in general, other neuromodulatory nuclei ^28^ may be at the higher end of correlation strength.

We detected synchronous activity among only 16% of cell pairs (~ 3000) and observed spike count correlation coefficients that were, on average, 0.04. Furthermore, strong noxious stimulation (electric foot shock) was expected to evoke a synchronized response of the entire LC population, but we instead observed that a relatively small proportion (~16%) of units contributed to the population response on each trial. Correlations decrease with lower firing rates ^44^ and urethane anesthesia is known to reduce spike rate by weakly effecting synaptic and non-synaptic currents ^40,65,79–81^. We report spike rates (mean 0.89 Hz), which were not far below what occurs during the awake state (reported means range from 0.92 Hz to 1.4 Hz in the rat and monkey ^2,45–50^). Moreover, when we considered correlations among only pairs with firing rates over 1 Hz, the proportion of correlated pairs remained similarly small (19.2%). Even more strikingly, similar levels of correlation were obtained for spontaneous and evoked firing (with the latter eliciting much higher spike rates), which further strengthens the view that LC synchrony may be much lower than previously assumed. Regardless of the anesthesia effects on the firing rate, many studies across many types of anesthetics, brain regions, and species have suggested that anesthesia actually increases spike count correlation coefficients by effecting large scale fluctuations of population activity ^39–42^. Thus, our finding of such low synchrony under anesthesia is unexpected. Building upon our findings, future work should test the hypothesis that, in the awake state and without the strong network fluctuations associated with urethane anesthesia or slow wave sleep, LC synchrony is further reduced.

The presumption about a robust LC synchrony originated from an earlier study, which estimated that the activity of 80% of LC cell pairs was synchronized over ~100 milliseconds; yet this estimation was based on a relatively small number (~20) of pairs recorded in the awake monkey ^32^. We also report that correlated activity was primarily focused in the timescale of less than 100 milliseconds based on spike train cross-correlograms assessed over a large range from 0.5 milliseconds to 40 seconds. Critically, however, the proportion of the population that was synchronized (16%) was much lower than the previous report (80%) when a larger population (>3000 pairs) was studied. Correlation strength is known to vary across brain regions and knowledge of the correlation structure in the LC lays the groundwork for understanding better the representations and computations of LC neurons and the functional role of noradrenergic neuromodulation by LC ensembles.

### Gap junctions as a mechanism underlying LC ensemble activity

Prior accounts have emphasized gap junctions as the source of synchrony in LC ^5,17,32^; however, we found little evidence to support this assumption. Numerous studies, which have inactivated gap junctions, conclusively demonstrate that the brief (0.5 to 1 millisecond) cross-correlogram peaks reflect gap junction coupling ^58–60^. In our data, we observed this sharp sub-millisecond cross-correlogram profile that may reflect putative gap junctions between LC neurons. However, the lack of sub-millisecond interactions beyond two neurons, as well as the rapid decay of sharp type interactions with distance each suggests that gap junctions are not likely to spread synchrony throughout the LC. It is unlikely that anesthesia interfered with observations of gap junction interactions. Although synchrony due to longer duration synaptic events (e.g., drive by common synaptic input to a cell pair) could be missed when spike counts are reduced under anesthesia (but see ^39–42^), synchrony due to brief events (e.g., driven by gap junctions) should not be affected by anesthesia-induced spike count suppression.

Intriguingly, it is possible that the gap junctions in the LC actually contribute to population desynchronization and the existence of simultaneously active LC ensembles. Systems that are coupled through gap junctions are certainly predisposed to exhibit synchronous changes of both membrane potential and spike probability ^82^. Recordings of the relatively deafferenated LC in slice (*in vitro*) recordings have demonstrated exactly such gap junction-synchronized membrane potentials ^17^. However, when whole-brain afferent input is present (as in our experiments) and the neurons have action potentials with a large amplitude after-hyperpolarization (as is the case for LC neurons ^30^), an excitatory afferent input can phase shift the relation between neurons’ membrane potentials, which results in both desynchronization at the population level as well as only sub-sets of neurons (ensembles) that are synchronized ^82^. Thus, our results do not support the view that gap junctions enable massive population synchrony in LC, but do not exclude their contribution to synchronize activity within distinct cell ensembles.

### Common synaptic inputs and local lateral inhibition may interact to promote LC synchrony

In addition to the sub-millisecond cross-correlogram peaks, we also observed cross-correlogram peaks over tens of milliseconds. These interactions over longer time scales may reflect a cell pair interaction with a third neuron (or group of neurons) that provides shared synaptic input to the pair. Intriguingly, we observed that these longer duration interactions were temporally diverse, spanning a range from 10 to 70 msec and lasting variable durations. While anesthesia could have caused under-sampling of the overall number of unit pairs synchronized by long duration synaptic events like common synaptic input, the existence of this temporal diversity of pairwise interactions is unlikely to be affected by anesthesia.

We propose that this delayed synchrony is due to a combination of shared synaptic inputs and lateral inhibition. A comparable time delay (±70 msec) has been previously reported in spike train cross-correlograms from paired intracellular slice recordings ^17^. In that work, the delayed cross-correlogram peaks did not occur when local LC activity was prevented, which is a manipulation that would remove local lateral inhibition. Moreover, the duration of lateral inhibition corresponds to this ±70 msec delay ^53^. Thus, our findings suggest the possibility that synchrony in the LC depends on the cooperation of shared synaptic inputs (extra-LC) and lateral inhibition (intra-LC). Wide type LC cells may be a crucial controller of LC synchrony given our findings that the wide type units may be the source of lateral inhibition.

### The mechanisms underlying the desynchronized population response to sensory stimuli

Our observations provide the first experimental evidence that LC single units do not spike synchronously in response to a noxious somatosensory input (foot shock). This result is surprising given that noxious sensory stimuli generate strong synaptic input across LC neurons, which is expected to drive correlated spiking among them. Noxious somatosensory stimuli are conveyed by both direct afferents from the dorsal horn neurons in the spinal cord (peripheral nociception) and the brainstem sensory trigeminal nuclei (peripheral nociception from the head) ^83^, as well as indirect (di-synaptic) nociceptive input from the rostral medulla ^84^. In both cases, single afferents synapse onto many LC neurons. We speculate that the desynchronized response to sensory stimulation may depend on each cell's pre-stimulus membrane potential, which may be in a different state prior to each sensory stimulus. We propose that this could occur given that LC neurons receive a diverse mixture of excitatory and inhibitory afferents from many brain regions – inputs that may be asynchronous (and/or opposing in excitatory/inhibitory valence) with respect to each other ^63,84,85^. Furthermore, refractory periods, self-inhibition, and lateral inhibition could set LC neurons to a dynamic mixture of membrane potentials. Thus, LC neurons that are in a higher state of excitability may spike and then inhibit other LC neurons from responding to the foot shock through lateral inhibition ^51,53,86^.

It is unlikely that sensory stimuli in other modalities, which only elicit a LC response in the awake state (e.g., non-noxious auditory, visual, and somatosensory stimuli ^49^), would evoke greater synchronous spiking than foot shocks do under anesthesia. Somatosensory stimulation that is milder than the noxious foot shocks we used here is not likely to evoke more robust LC responses in awake state. It is possible that stimulation that is close to the physiological limit may increase the probability of a synchronized population response ^87^. Visual and auditory stimuli evoke synaptic responses in the LC via a longer poly-synaptic pathway ^88–90^. The synaptic delays and the noise added by synaptic transmission at each step are expected to jitter the timing of common synaptic input to LC cell pairs and, thus, reduce synchronized spiking. These questions should be resolved by recording large numbers of single units from the awake animals.

### Potential for targeted neuromodulation by LC ensembles

Contrary to the current view of the LC as a non-specific neuromodulatory system, our data suggest that targeted forebrain neuromodulation could possibly be achieved by selective activation of gap junction coupled LC cell assemblies that share common efferent targets. We showed that unit pairs with synchronous activity on the sub-millisecond timescale were more likely to project to similar forebrain regions. We also observed a tendency for anti-correlated units, which spike in opposition with one another, to avoid projecting to the same forebrain areas. These data are consistent with targeted neuromodulation under a scenario in which one population of units projects heavily to the forebrain Region A, while another population avoids Region A and projects heavily to Region B. During times when the A-projecting population is active, NE content in Region A increases while the B-projecting population is suppressed and NE content in Region B decreases.

Targeted forebrain neuromodulation may also be achieved by NE gradients between forebrain sites, which could be generated by infra-slow (<1 Hz) oscillations in spike rate. Our results demonstrate that LC neurons will have in-phase spike rate oscillations within their group and anti-phase spike rate oscillations with neurons outside their group. Thus, over an infra-slow duration (2 to 10 sec), some LC neurons will synchronously release NE to their projection targets, while other LC neurons are suppressed. Depending on their projection targets (which can often be single forebrain regions according to our data and others ^29^), NE could be simultaneously high in some forebrain regions and simultaneously low in others over a timescale of 2 to 10 sec.

We also speculate that these infra-slow changes in LC spike rate could orchestrate infra-slow LFP, EEG, or BOLD oscillations in multiple cortical regions and promote communication among these regions by synchronizing their infra-slow oscillations in order to organize functional (or resting state) networks ^67–70,91,92^. LC spiking (and associated NE release) regulates cortical excitability; coherent infra-slow oscillations of LC spike rate among sub-sets of LC neurons may therefore allow different LC ensembles to influence synchronization among the regions in separate cortical networks. Our data thus provide experimental support for a theoretically predicted, but speculative function of the LC to organize task-related and resting state networks ^67,93,94^. Given that different LC cell types spiked coherently at different infra-slow frequencies, it is possible that narrow and wide cell types participate differentially in resetting cortical networks. Additionally, vascular innervation by LC neurons may directly affect the hemodynamic response and alter the physiology underlying the generation of the fMRI BOLD signal used to study cortical networks. The physiology of the BOLD signal, which is a common research and diagnostic tool, may be better understood when future studies establish how LC neurons – and especially the different LC cell types reported here – modulate cortical network-specific neurovascular coupling.

Lastly, our findings do not contradict the long-standing notion of global NE neuromodulation. Volume NE release may provide simultaneous post-synaptic neurotransmission in distant brain regions on a time scale of a few seconds ^11,95^. This global and relatively slow neuromodulation is clearly synchronous by nature and presents a critical component of controlling brain state and neuronal excitability, especially on behaviorally relevant timescales (seconds, minutes, or hours). In addition to its role in global neuromodulation, the LC system appears to be anatomically and functionally differentiated with diverse cell types and finely-structured activity patterns. These features may allow a more nuanced role for the LC in theories of systems and cognitive neuroscience.

## Methods

### Animals

Twelve male Sprague-Dawley rats (350 - 450 g) were used. All experimental procedures were carried out with approval from the local authorities and in compliance with the German Law for the Protection of Animals in experimental research (Tierschutzversuchstierverordnung) and the European Community Guidelines for the Care and Use of Laboratory Animals (EU Directive 2010/63/EU).

### Anesthesia and Surgical Procedures

Rats were anesthetized using an intra-peritoneal (i.p.) injection of a 1.5 g/kg body weight dose of urethane (Sigma-Aldrich, U2500). Oxygen was administered. The animal was placed on a heating pad and a rectal probe was used to maintain a body temperature of 37 C. The eyes were covered in ointment. After removal of the skin, the skull was leveled to 0 degrees, such that the difference between lambda and bregma was less than 0.2 mm.

### Stereotaxic coordinates and electrode placement

Craniotomies were made at the locations listed in the following table (Extended Data Table 1). Accurate electrode placement was confirmed by examining the firing properties of neurons in each brain region. In the LC, these criteria included a slow spontaneous firing rate, biphasic response to noxious sensory stimuli (foot shock), audible presence of jaw movement-responsive cells in the MeV (Mesencephalic Nucleus of Cranial Nerve V) with undetectable single units (<0.2 mV). LC electrode placements were later verified using histological examination of brain tissue sections (Extended Data Figure 8). Placement of electrodes for forebrain stimulation was stereotactically-guided and, when possible, electrophysiological criteria were used to verify target placement.

### Electrodes

Stimulation of cortical and sub-cortical brain regions was conducted via tungsten electrodes with low impedance (10 - 50 kOhm) to prevent heating at the electrode tip. Tungsten electrodes with a diameter of 200 μm (FHC, Model: UEWMFGSMCNNG) were ground at the tip to lower impedance to this range. Recording from the LC used a 15 μm thick silicone probe with 32 channels (NeuroNexus, Model: A1x32-Poly3-10mm-25s-177-A32). The channels were implanted toward the anterior aspect of the brain. Each channel was separated from the neighboring channels by 25 μm. Channels were divided into 10 tetrodes with one channel overlapping per tetrode (Extended Data Figure 1). The 275 μm extent of the recording channels covered nearly the entire dorsal-ventral extent of the LC, which is ~ 300 μm ^96,97^.

### Recording and signal acquisition

A silver wire inserted into the neck muscle was used as a reference for the electrodes. Electrodes were connected to a pre-amplifier (in-house constructed) via low noise cables. Analog signals were amplified (by 2000 for LC and 500 for cortex) and filtered (8 kHz low pass, DC high pass) using an Alpha-Omega multi-channel processor (Alpha-Omega, Model: MPC Plus). Signals were then digitized at 24 kHz using a data acquisition device CED, Model: Power1401mkII). These signals were stored using Spike2 software (CED).

### Spike detection

The recorded signal for each channel was filtered offline with a four pole butterworth band pass (300 - 8000 Hz). Spikes were then detected as crossings of a negative threshold that was four times the standard deviation of the channel noise. Noise was defined as the median of the rectified signal divided by 0.6745 ^98^. Detected spike waveforms were stored from 0.6 msec to 1.0 msec around the threshold crossing. This duration was chosen based on the known action potential duration of rat LC neurons ^2,99^. A 0.6 msec refractory period was used to not detect a subsequent spike during this window.

### Spike clustering

Spike waveforms were clustered using an automatic clustering algorithm and then manually refined and verified using cluster visualization software (CED, Spike 2). Automated clustering was performed using Wave_Clus ^98^ in MATLAB (default parameters for clustering) followed by manual refinement and verification in clustering visualization software. This method uses wavelets to decompose the waveform into a simpler approximation of its shape at different frequencies (wavelet scales) and times in the waveform. Using this method, small amplitude bumps or deflections at different time points in the waveform can be used to cluster waveforms together, if they are a highly informative waveform characteristic. After the automated sorting, manual refinement of clustering using a 3-dimensional plot of principle components or the amplitude at particular waveform time points (peaks and troughs). Auto-correlograms were used to assess the level of noise (refractory period violations) and cross-correlograms between simultaneously recorded units were used to prevent over-clustering ^100^.

### Detection of spikes across tetrodes

Due to configuration of the recording array with a high-density of electrode contacts, the spikes from the same LC neuron could be detected on more than one tetrode. In such situations, we first attempted to merge the spike trains across multiple tetrodes. Merging the spike trains potentially originating from the same neuron and detected on multiple tetrodes should reduce false negatives (missing spikes), as it is common for PCA of waveforms to miss some spikes even if they are part of a well isolated cluster. The assumption is that different spikes are missed on different tetrodes, which were subjected to separate PCA's during clustering. Therefore, spikes missed by PCA on one tetrode could be partly filled in by spikes that were detected on other tetrodes, providing that the tetrodes were recording the same single unit. Merging spike trains across tetrodes operated on the principle that, if the spikes from the same neuron are recorded, for example, on 3 adjacent tetrodes and the spike waveforms can be classified into 3 well-isolated clusters, then the spike trains can be merged to yield an equally well-isolated unit. Furthermore, merging across spike trains allows units to be tracked if they drift away from one tetrode and become closer to another tetrode. However, the merging procedure needs to avoid inclusion of contaminated spikes originating from neighboring cells. Therefore, merged spike trains must be statistically tested for false positives and conservatively discarded. Unit cluster contamination from other units are typically detected by the presence of spikes during the refractory period. Thus, if the merged spike train did not meet criteria for a single unit activity, then we kept the unit recorded on the tetrode with the least noise (lowest proportion of spikes during refractory period).

The merging process consisted in the following steps. First, the cross-correlograms were computed at the sampling rate of the recording (0.04 ms bin width) between the spike trains of a cluster isolated from one tetrode (“reference” cluster) and all other clusters isolated from all remaining tetrodes. If the spike trains associated with the two clusters contained spikes from the same unit, then the majority of spikes would have identical timing with the vast majority (>90%) of spikes being coincidental at time 0 (with a few sampling points of error) and the remaining spikes being spikes either detected on one tetrode and missed on the other or cluster noise from other units. Prior recordings using high-density linear electrodes have used cross-correlograms to simply discard one of the trains ^101^; however, we used merging across tetrodes to reduce missed spikes. In the case that the "reference" and "other" spike trains were mostly coincidental spikes, we attempted to merge them using a procedure, as follows. The coincident spikes in the reference spike train were deleted and the remaining reference spikes were merged with the spikes from the other spike train, resulting in a new "merged" spike train. The amount of noise (number of spikes during the refractory period) and total number of spikes in the train were recorded for the original "reference" spike train, original "other" spike train, and newly merged spike train. A Fisher’s Exact Test was performed to statistically assess if the proportion of contaminated spikes added by merging is significantly greater than the proportion of noise (refractory period) spikes in either the original reference spike train or the original other spike train. The Fisher's test was run separately for merged versus reference and the merged versus other spike trains. A result of non-significance (with alpha set to 0.01 and a right-sided test) for *both* the original reference spike train and the original other spike train indicated that the original spike trains do *not* have *greater* odds of having noise than the merged spike train. In this case, the merged train may be kept because the amount of noise was not increased beyond its level in the original two trains. The original spike trains were discarded. However, if the test is significant for either of the original trains, then either the reference spike train or the other spike train is kept depending on which has a lower percentage of spikes during the refractory period out of the total number of spikes. The merged spike train was discarded. The merging procedure was repeated for each cluster from each tetrode until conflicts no longer existed.

### Characterization of LC units

Single units were identified using typical criteria (Extended Data Figure 1). These criteria were low firing rate (0.89 spikes/sec) and a characteristic bi-phasic response (excitation followed by inhibition) to sensory stimuli. The neurochemical nature of LC units was identified by the presence of auto-inhibitory alpha-2 adrenergic receptors which were activated using the alpha-2 agonist, clonidine. Electrode tracks approaching LC were visualized (Extended Data Figure 8). These methods are the standard for demonstrating that the neurons are likely to be LC-NE neurons. Therefore, our results are comparable with the existing literature on the LC and we were likely sampling mostly LC-NE neurons.

### Assigning unit location on recording array

Isolated single units were assigned a channel location on the electrode array according to which electrode measured the highest mean waveform amplitude (averaged from all spikes). In the case of single units with spikes that were merged across tetrodes, a list tracked the tetrodes on which the unit appeared and the location was assigned to the channel with the maximum amplitude when considering all of the tetrodes that recorded the unit. The spacing between all 32 channels was 25 μm, which allowed us to use Pythagoras’ Theorem to calculate a distance between channels on the array. The distance between each unit's maximal waveform amplitude was used to measure the distance dependence of spike count correlations and cross-correlograms.

We inferred the spatial probability distribution of narrow and wide units on the array by fitting N^th^-order polynomials (N = 2 to 9) to the proportion of each unit type recorded at each of the 10 tetrodes. We found that N = 5 or 6 produced an R^2^ that was > 0.9, whereas lower N had poor fits (R^2^=0.3 to 0.7) and larger N visually overfit the data. The polynomial function provided a probability, which we verified in all rats by removing one rat and re-calculating the fit until all rats had been removed once (jackknife error). In all cases, except one, R^2^ was >0.9.

### Sensory stimuli

Sensory stimuli were foot shocks delivered to the contralateral hind paw. Pulses were square, biphasic, and 0.005 msec duration at 5 mA. Pulses were delivered at two frequencies (single pulse or five pulses at 30 Hz) delivered in random order. Fifty trials of foot shock stimuli were delivered with an inter-trial interval of 2000±500 msec.

### Intra-cranial stimulation

Brain regions were stimulated in a random order. Pulses were square, biphasic, and 0.25 msec duration over a range of intensities (400 - 1200 µA), which were delivered in a randomized order. The pulse waveforms were constructed in Spike2 (CED) and delivered via a current generator (in-house constructed), which allowed recording of the stimulation voltage at the tip of electrode, which was also digitized and stored and used to verify stimulation. Stimulation was delivered with an inter-trial interval of 2000±500 msec. At least 50 trials of each intensity were delivered for each brain region.

### Administration of clonidine

At the end of the recording, a 0.5 mg/kg dose of the alpha-2 adrenergic agonist clonidine was injected i.p. (Sigma-Aldrich, product identification: C7897). The recording was continued at least until all activity, included multi-unit activity, ceased.

### Histology

Rats were euthanized with pentobarbital sodium (Narcoren, Merial) via an i.p. injection (100 mg/kg). The rats were then trans-cardially perfused with 0.9% saline and then 4% paraformaldehyde (PAF) in 0.1M phosphate buffer (pH 7.4). The brain was removed and stored in 4% PAF. Brains were moved into 30% sucrose, until they sank, before sectioning on a freezing microtome (Microm, model: HM 440E). Coronal sections (50 μm thick) were collected into 0.1M phosphate buffer and directly stained. For sections containing the LC, alternating sections were stained for Nissl substance or the catecholamine synthesis enzyme, tyrosine hydroxylase (TH). Sections containing cortical and sub-cortical regions were stained for Nissl substance. Staining for TH was performed using a 1:4000 dilution of monoclonal mouse anti-TH antibody (ImmunoStar) and a 1:400 dilution of biotinylated, rat absorbed anti-mouse secondary antibody (Biozol) in PB. The antibody was visualized using a DAB and horse radish peroxidase reaction with hydrogen peroxide using a standard kit (Biozol, model: Vectastain Elite ABC-Peroxidase Kit CE). After staining for TH or Nissl, sections were mounted on gelatin-coated glass slides. Nissl stained and slide mounted sections were dehydrated in an alcohol series. Slides were cleared (Carl Roth, Roti-histol) and cover slipped with DPX slide mounting media (Sigma-Aldrich, catalog number: 06522).

### Data analysis: Spike count correlations

The Pearson’s correlation coefficient was used to quantify the correlation between spike counts. Spontaneous spiking excluded the 1 sec period following foot shock stimulation or intra-cranial stimulation. Spontaneous spikes were also excluded during inactive periods in which the rate was less than 0.5 Hz due to quiescence of all single and multi-unit activity in the LC (paradoxical sleep ^2^). Spontaneous spike count correlations were then calculated from the time bins (200 msec or 1000 msec) in which both neurons were active.

Poisson spike trains should generate some degree of synchrony (spike count correlation coefficient) by chance. We compared the correlation coefficient for each pair against 500 surrogate spike trains for the same pair. The trains were generated in manner that preserved the inter-spike interval (ISI) distribution for each unit and the slow (<2 Hz) oscillations in spike rate (thought to be generated by brain network-wide interactions under anesthesia). Each surrogate spike train had the same oscillation phase preference, the same number of spikes per oscillation cycle, and an identical ISI distribution, but the precise spike timing of each unit in relation to other units is randomized.

Evoked spike count correlations after foot shocks were calculated from the trial-by-trial spike count in a single window after stimulus onset (50 msec). This window was chosen based upon the timing of the spiking evoked by a single pulse foot shock in our recordings, as well as reports by others ^16,49^. For evoked correlations, the spiking on each trial was sparse in the 50 msec window, so a statistical permutation test was under-powered and could not be performed.

### Data analysis: Cross-correlograms

We calculated cross-correlograms between spike trains. Significant changes in coincidental spike count were detected by comparing the observed counts to 1000 surrogate cross-correlograms generated from jitter spike times ^102^. This approach uses the data to determine the degree of coincident spiking expected by chance and it also excludes synchrony due to interactions at slower time scales than those of interest. Briefly, the spike times for each unit were jittered on a uniform interval and a surrogate cross-correlogram was calculated between the jittered spike times; this process was repeated 1000 times. Significant cross-correlogram bins were those that crossed both a 1% pairwise expected spike count band and a 1% globally expected spike count band (the maximum and minimum counts at any time point in the cross-correlogram and any surrogate cross-correlogram). For broad-type interactions, the cross-correlograms were calculated in a window of 2000 msec, a bin size of 10 msec, and a uniform jitter interval of ±200 msec. Any significant coincidental spiking excludes synchrony due to co-variation of spiking on a timescale of a few hundred milliseconds. For sharp-type interactions, we used a window of 3 msec, a bin size of 0.05 msec, and a uniform jitter interval of ±1 msec.

To ascertain if we missed interactions at other timescales, we employed the method of integrating the cross-correlogram ^36^. The gradual integration of the cross-correlogram in successively larger windows (e.g., τ= ±5 msec, 10 msec,…40,000 msec) will result in a curve that changes rapidly during tau with a large concentration of coincidental spiking and will eventually plateaus at very large tau at which the firing patterns of the pair is unrelated. We integrated in 1 msec steps from 0 to 40 sec and recorded the start of the plateau. The results of this analysis suggested that some interactions could occur on the level of a few seconds. Thus, we calculated additional cross-correlograms using a window of 20 sec, a bin size of 0.2 sec, and a uniform jitter on the interval of ±2 sec.

### Data analysis: Syn-fire chain analysis

Repeating sequences of triplets of neuronal spiking were measured using three steps: (*i*) a spike-by-spike search, (*ii*) template-formation, and (*iii*) template matching algorithm ^103,104^. The analysis stepped through all simultaneously recorded units *n*→*N* and all of the spike times (*M*) for each unit (*n*_*m*→*M*_). The spike search started with the first unit and its first spike, *n*_*m*_. This spike time was a reference event marking the start of a 2 msec window during which we identified any other spiking units. The sequence of units and the delay between their spikes was stored as a template. For example, *n*_*m*_ might be followed 1.1 msec later by a spike from unit *n* = 5 which is subsequently followed 0.8 msec later by a spike from unit *n* = 30. This forms a template of 1 – 5 – 30 with delays of 1.1 msec and 0.8 msec. The next step, template matching, would proceed through the spikes 1_*m*+1→*M*_ and attempt to match the template and its delays. A 0.4 msec window of error was allowed around each spike in the original template for matching. Thus, for each spike of unit *n* = 1, unit *n* = 5 could spike between 0.7 msec and 1.5 msec after unit *n* = 1 and unit *n* = 30 could spike 0.4 msec to 1.2 msec after unit *n* = 5. A template match would be counted if the spikes of the other units aligned with the originally formed template. For each template, the total number of observations was compared to the number of observations in the top 1% of 100 surrogate data sets in which spike times were jittered on a uniform interval by 1 msec. Any sequential spike patterns that occurred more often than expected by chance were counted as significantly occurring chains of spikes.

### Data analysis: Graph theory analysis and ensemble detection

For each rat, a graph was constructed with each unit as a node. Links were drawn between units with strong spike count correlations, following the methods of Rubinov & Sporns (2010) and Bruno et al. (2015) ^105,106^. The threshold for drawing a link was set as the highest possible value such that the mean network degree was less than the log(N), where N is the number of nodes in the graph ^107^. Units without strongly correlated activity were left unlinked. The resulting network was represented by a binary adjacency matrix. Ensembles were detected by segregating nodes into groups that maximize the number of links within each group and minimizes the number of links between group. The iterative optimization procedure used a Louvain community detection algorithm to maximize a modularity score (Q) quantifying the degree to which groups separate ^106,108^. The degree of ensemble separation (Q) was compared to modularity scores from 1000 from shuffled networks. If Q was higher than the top 5% of the 1000 surrogate values, then adequate separation of units into ensembles was achieved. All graph theory and ensemble detection analyses were implemented in MATLAB using the Brain Connectivity Toolbox ^109^.

### Data analysis: Oscillations in spike count

Single unit spike trains were first convolved with a Gaussian kernel with a width of 250 msec and a sampling rate of 1 msec. The resulting spike density functions (SDF) were analyzed for the power of oscillations, the phase of oscillations, and the coherence between pairs of SDF’s. The power spectral density of each spike train was calculated using a multi-taper method in MATLAB (Chronux Toolbox) ^110^. We used 19 tapers with a time-bandwidth product of 10. The frequency band was 0.01 to 10 Hz. We used finite size correction for low spike counts. Instantaneous phase was extracted by first filtering the SDF with a 3^rd^ order Butterworth filter at a particular frequency of interest (0.09 to 0.11 Hz and 0.40 to 0.48 Hz) and then obtaining phase from the Hilbert transform of the signal. The consistency of the instantaneous phase difference between each unit pair was assessed using Rayleigh’s Test for Circular Uniformity (p<0.05), which was implemented in MATLAB (CircStat Toolbox) ^111^. Coherence between pairs of units was calculated using the Chronux Toolbox in MATLAB with the same parameters used for calculating the power spectral density. Coherence and power were averaged across all single units and smoothed with a width of 0.15 Hz.

### Data analysis: Spike-LFP phase locking

Local field potential (LFP) was recorded in the prefrontal cortex. The LFP was lowpass filtered with a 3^rd^ order Butterworth filter at 2 Hz. The instantaneous phase was obtained by Hilbert transformation. Spike times corresponded to LFP phases. For each single unit, the phase distribution was tested for significant locking to cortical LFP using Rayleigh’s Test for Circular Uniformity (p<0.05).

### Data analysis: Antidromic spiking

Forebrain regions were stimulated in a randomized order with single pulses at currents of 400, 600, 800, 1000, and 1200 µA. Stimulation was delivered with a 2 sec inter-trial interval with 500 msec jitter. Peri-stimulus time histograms were Z-scored to 1 sec before stimulus onset. If a single bin (5 msec), but no other bins, exceeded a Z-score of 5, then the unit was marked as antidromically activated. The bin size was chosen, based on prior work, which has demonstrated that 3 or 4 msec of jitter occurs when stimulating LC fibers because they lack myelin ^112^. This rationale is based on an extremely low chance of consistent spiking within the same 5 msec window by slowly firing neurons (typical ISI is longer than 100 msec). However, manual inspection of the individual spike rasters was used to confirm the results.

### Data analysis: statistics

Data were tested for normality using a Shapiro-Wilk test (alpha = 0.05) and homogeneity of variance (alpha = 0.05) using an F-test (vartest2 in MATLAB) for 2 groups or Levene’s Test (vartestn in MATLAB) for more than 2 groups. If data were not normal, then a Wilcoxon-Mann-Whitney Test was used for 2 groups or a Kruskal-Wallis Test for more than 2 groups; otherwise, a two-sided t-test or between-subjects one-way ANOVA was used. If variance was inhomogeneous, then we used Welch’s t-test or Welch’s ANOVA. For ANOVA (or Kruskal-Wallis), post-hoc testing between individual groups was performed using the MATLAB function, multcompare (with Dunn-Sidak correction for Kruskal-Wallis). If a Welch’s ANOVA was used for heteroscedastic data, then post-hoc testing was performed using the Games-Howell test. These comparisons were unplanned. Mean and standard error are reported for normally distributed data. Median and standard deviation are reported for data that were not normally distributed.

We report effect sizes as Cohen’s D for analysis of 2 groups (e.g., t-test or Wilcoxon-Mann-Whitney) or, for analysis of more than 2 groups, we report ω^2^ (e.g., ANOVA) or ω^2^ (e.g., Welch’s ANOVA). For 2 × 2 tables, we used a Fisher’s Exact Test, for which the effect size is quantified by the Odds Ratio (OR), which we converted to Cohen’s D (termed D_or_) ^113^.

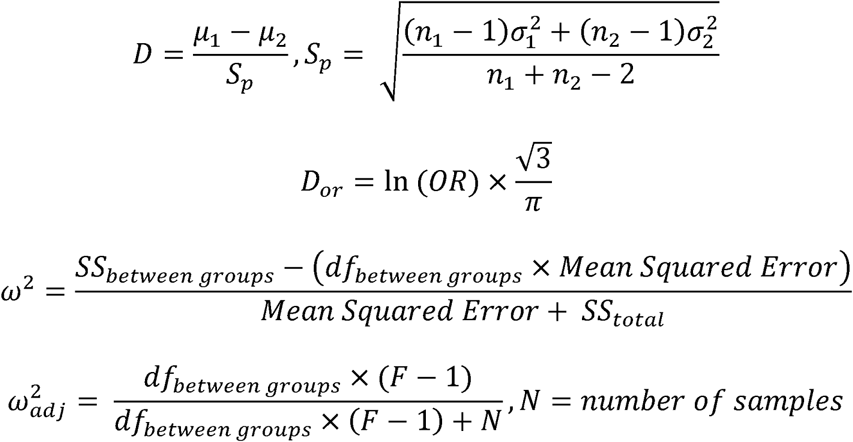

In the case that a null hypothesis was rejected, we made a post-hoc determination that the sample size and statistical test provided adequate power to reject the null hypothesis. The power was calculated with sampsizepwr in MATLAB for 2 groups. If there were more than 3 groups, then the power was calculated using powerAOVI in MATLAB. We are unaware of methods for assessing the power of a Kruskal-Wallis Test or a Welch’s ANOVA and do not report power for those tests.

For circular data, uniformity was assessed using Rayleigh’s Test for Circular Uniformity (alpha = 0.05). These calculations were made using the CircStat toolbox in MATLAB ^111^. Power calculations were not made for circular statistics.

### Data availability statement

The generated datasets for this study are available from the corresponding author on reasonable request.

### Code availability

The code used for data analysis are available from the corresponding author.

## Acknowledgments

We thank Axel Oeltermann and Eduard Krampe for technical assistance with stimulating electrodes, Joachim Werner for technical support, Dr. Henry Evrard for expertise in immunohistochemistry, Marcel Hertl, Felicitas Horn, Jennifer Smuda, and Katalin Kalya for assistance with histology, and Valeria d’Andrea for helping to write the code for the permutation statistical test. We thank Dr. Susan Sara for comments on the paper. This research was supported by the European Union’s Marie Curie Fellowship in the FP7 funding scheme to N.K.T. (PIIF-GA-2012-331122) and the European Union’s FET Open in the FP7 funding scheme (SICODE) to S.P., O.E., and N.K.L. All data are archived for 10 years or longer at the Max Planck Institute for Biological Cybernetics. The author contributions are as follows: N.K.T., N.K.L., O.E. - conceived of the project; N.K.T., O.E. – experimental design; N.K.T., S.P., N.K.L., O.E. – analysis design; N.K.T., R.N. - collected data; N.K.T. - analyzed data; N.K.T., O.E. - wrote the paper; all authors - edited the paper.

## Extended Data

**Extended Data Figure 1.**
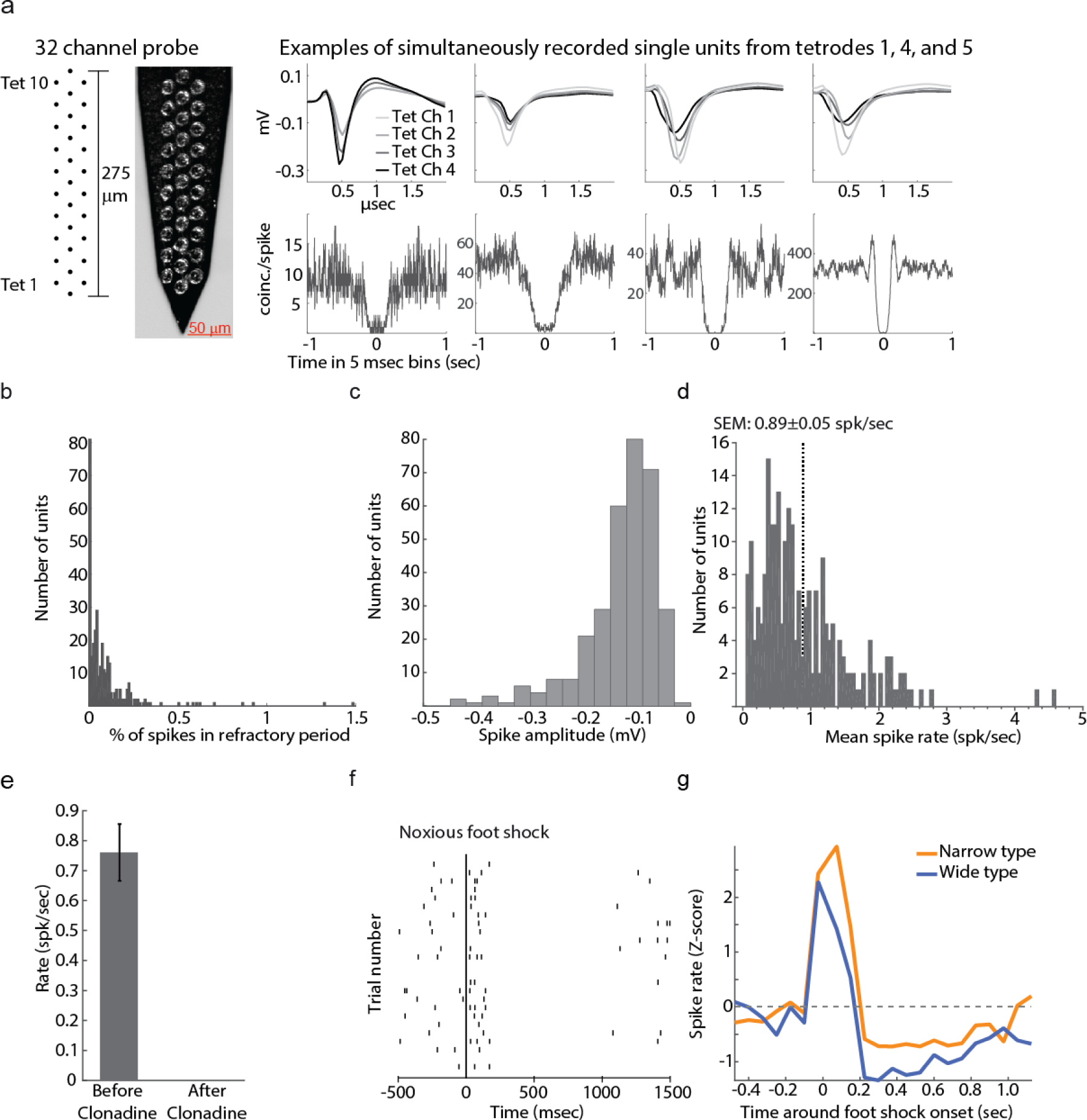
Characterization of single unit spontaneous activity, response to sensory stimuli, and clonidine. A. Units were recorded using a silicone shank with 10 tetrodes. Each tetrode contained 4 channels with one channel overlapping with the adjacent channel. The probe was advanced until units on all channels were responsive to foot shock and inhibited by clonidine. The rightward panel shows the average waveforms recorded on each tetrode channel for 4 units recorded on 3 different tetrodes. Below the waveforms, the auto-correlograms show a typical inter-spike interval of > 100 msec. B. A histogram showing the distribution of refractory period violations (set at a conservative limit of 10 msec) across all units. Overall, the proportion of spikes during the refractory period was < 2% with most neurons having fewer than 0.25% of their spikes during the refractory period. C. A distribution of single unit spike amplitudes taken from the tetrode channel with the maximal waveform. Spikes were, in general, of high amplitudes allowing reliable detection and sorting of spikes. D. The distribution of single unit firing rates shows that firing was typical of the LC with a mean of 0.89 spikes per sec. E. Units were inhibited by clonidine. The average firing rate before and after clonidine is plotted. F. An example raster showing the biphasic response to foot shocks (at t=0 msec). Each row is a trial and the ticks represent spikes. G. The normalized mean response profile for narrow and wide units is plotted around foot shock onset at time 0. This plot illustrates the response to a burst of foot shocks (5mA pulses delivered at 30 Hz).

**Extended Data Figure 2.**
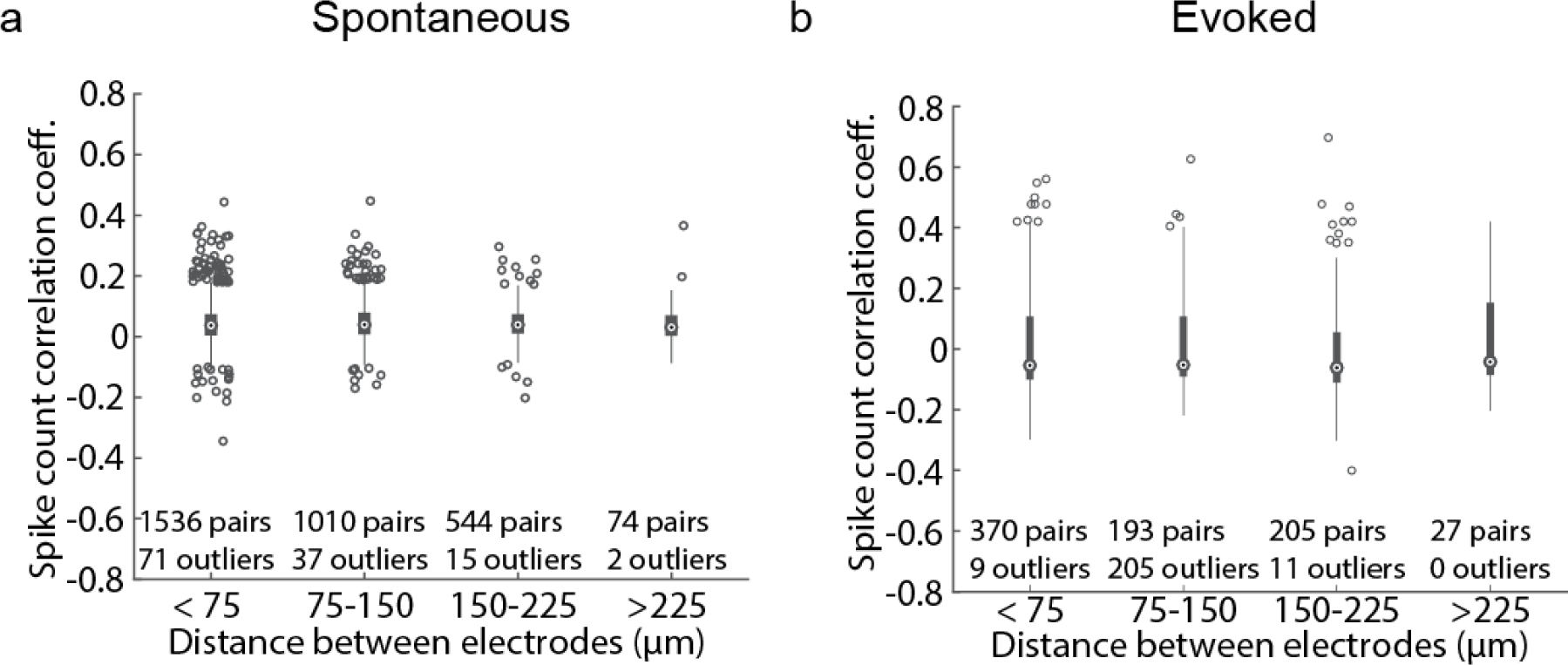
Spike count correlation coefficients did not depend on distance between unit pairs. The distance between units was estimated as the distance between the electrode contacts that recorded the maximal amplitude of each unit. Data are plotted as box plots.

**Extended Data Figure 3.**
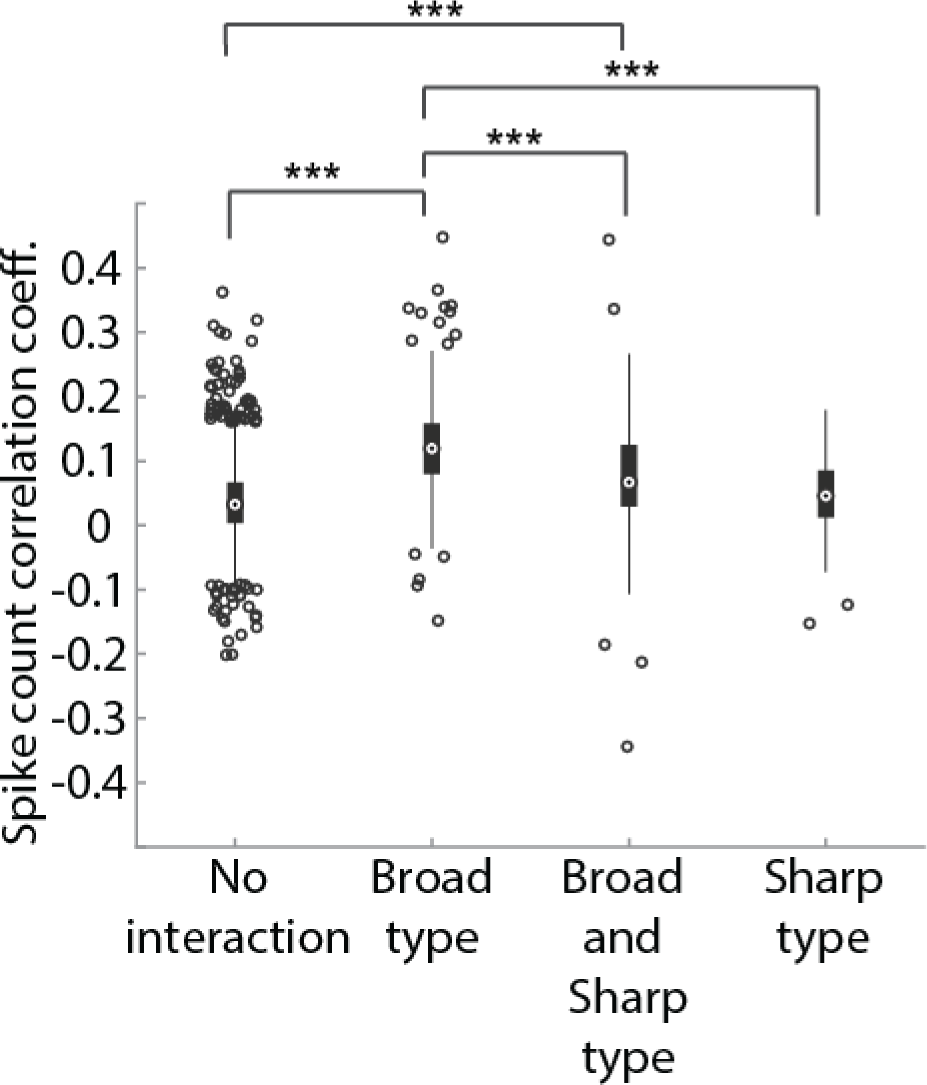
Pairs with network interactions have higher spike count correlations.

**Extended Data Figure 4.**
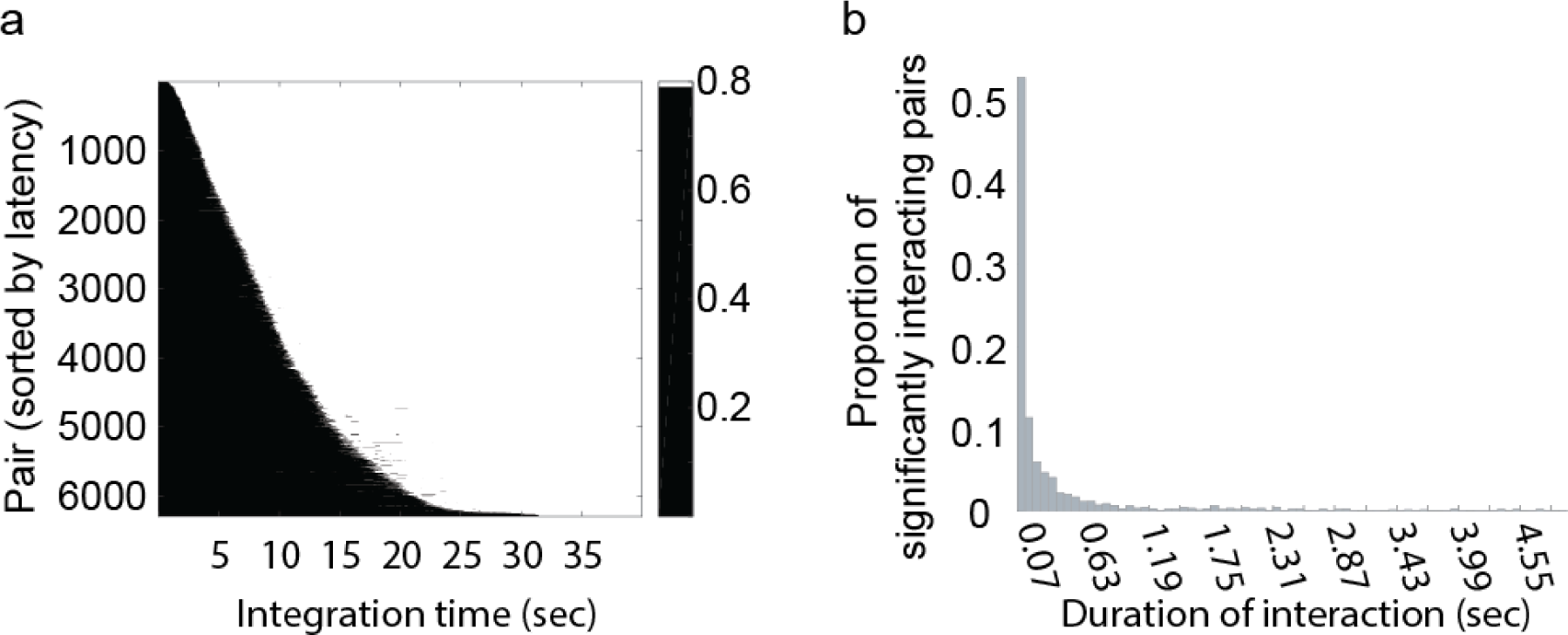
Synchrony in spike train cross-correlograms over the time scale of seconds was extremely rare. (A) The cumulative correlation coefficient was obtained at various tau by integrating over successively larger windows of the spike train cross-correlograms calculated over a ±40 sec window in 5 msec bins. For each recorded pairwise cross-correlogram (considered in both directions) on the y-axis, the value of the cumulative correlation coefficient (black-white color) is plotted against tau (in seconds, x-axis). The tau at which the integration saturates is approximated at 0.8 (white). This point estimates when the majority of an interaction (a bump on a cross-correlogram) has ended and thus gives an overview of the timescale of interactions present in the data. This analysis indicated that interactions may occur up to 20 sec. (B) In order to test for interactions between 20 msec and 20 sec in duration, we calculated cross-correlograms and measured the duration of the interaction (excess coincidental spikes beyond the 1% pairwise global confidence interval derived from a surrogate sets of ±2 sec jittered spike trains). The analysis revealed that the majority of significant interactions have a duration of under 1 sec with many lasting approximately 70 msec.

**Extended Data Figure 5.**
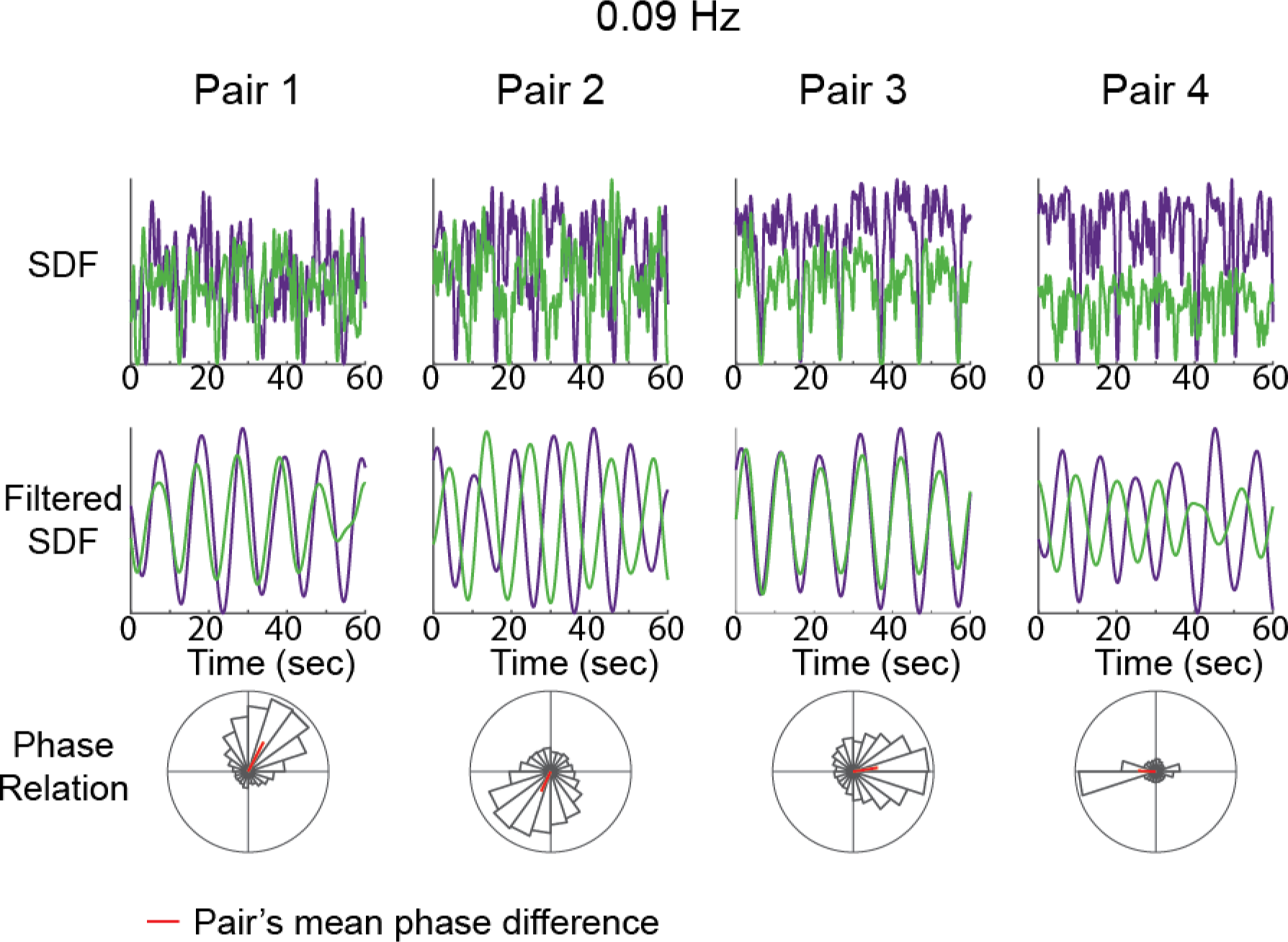
Examples of pairwise spike rate oscillations at 0.09 Hz. The top and middle panels shows spike density functions over a small recording segment; the bottom panels show the phase relation between the units in the pair over the entire session. The mean phase relation for each pair is marked by the red line. All example pairs had a significantly non-uniform (Rayleigh’s Test for Circular Uniformity, p<0.05).

**Extended Data Figure 6.**
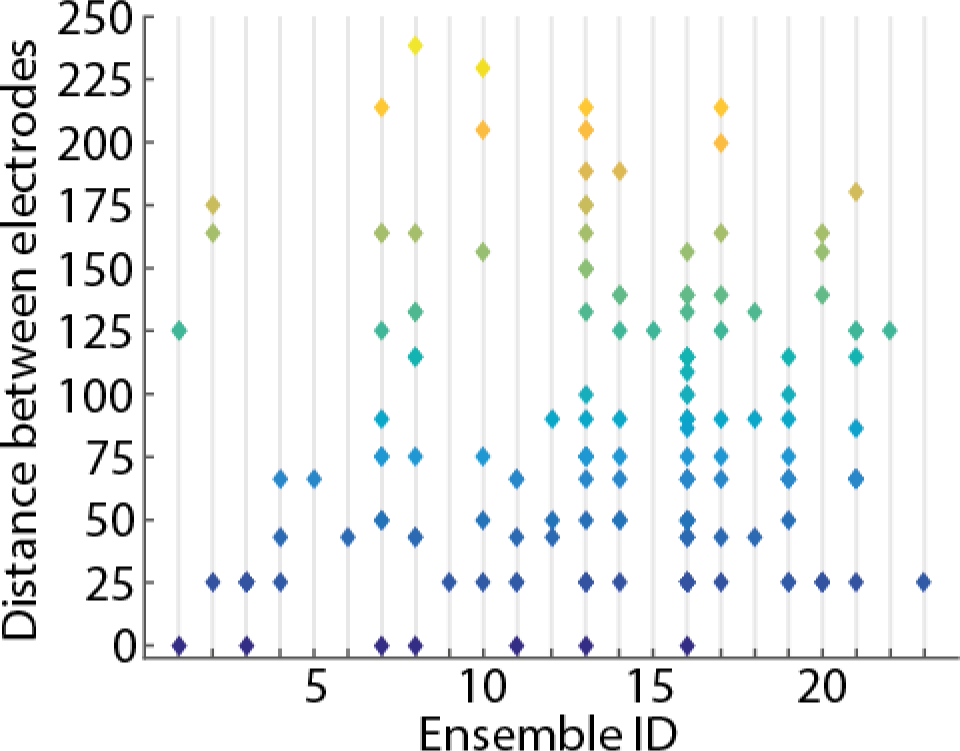
Ensembles are spatially diffuse. The pairwise distance between all pairs within an ensemble are plotted for all 23 ensembles. The y-axis and the color indicate the distance between all unit pairs in each ensemble.

**Extended Data Figure 7.**
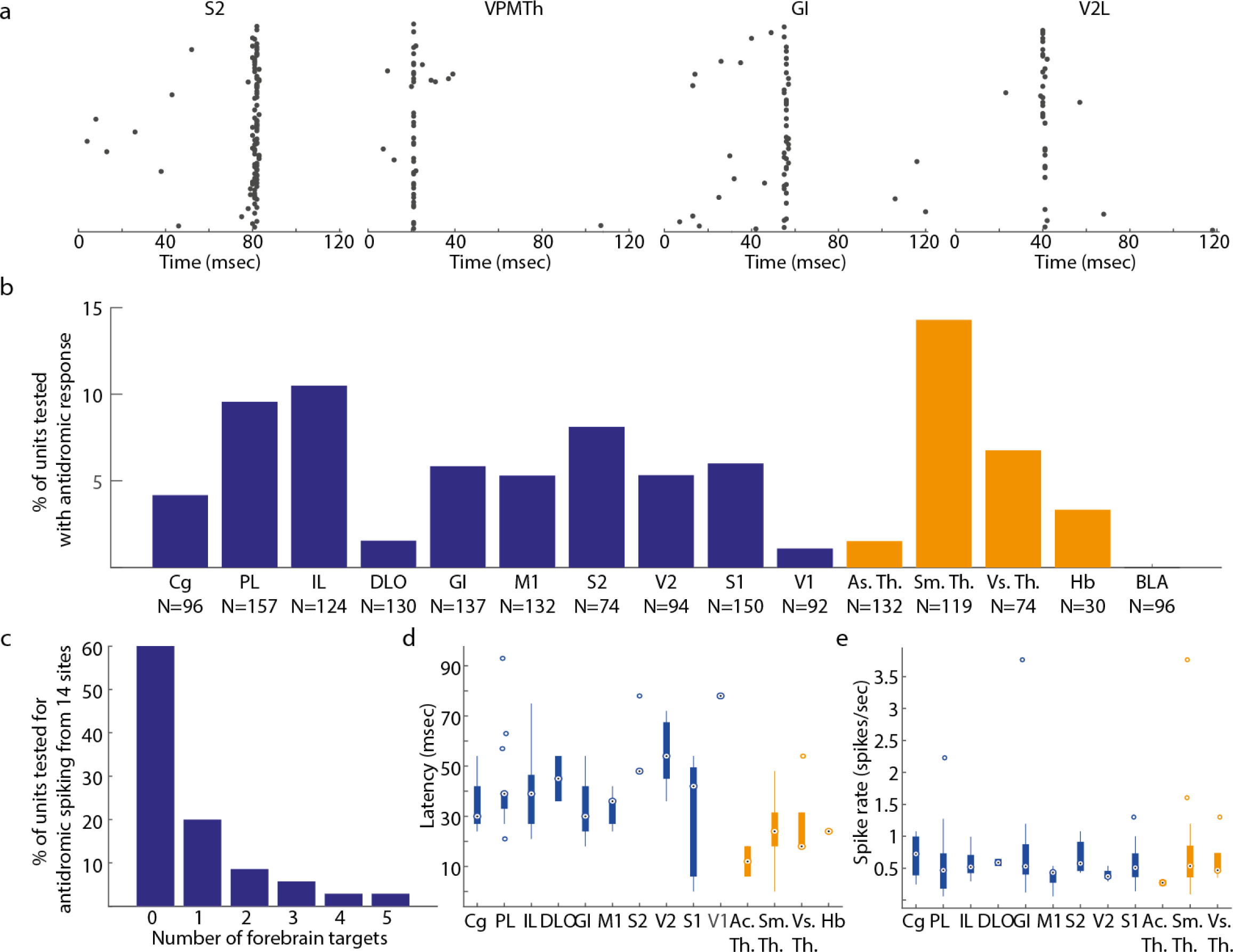
A summary of forebrain projection patterns and latencies. A) Examples spike rasters showing the timing of antidromically-drive spikes. The jitter of a few milliseconds and lack of consistent response on every trial is typical of unmyelinated LC axons ^112^. (B) Single units projected to a variety of forebrain sites. The y-axis shows the percent of units projecting to each site. The total number of units tested for projections to each site is written on the x-axis. Cortical regions are in blue and sub-cortical regions are in orange. (C) A mixture of broad (multiple targets) and selective (single target out of 15 regions tested) projection patterns were observed. Antidromic activation of units after stimulation ranged from 1 to 5 forebrain sites. Selective projections are in agreement with prior anatomical studies that traced projections of single LC neurons ^29^. The average number of projections per single unit with antidromic spiking was 2.0±0.3, which is similar to the 1.6±0.8 projection targets reported using barcoded RNA ^29^. (D) A box plot shows the latency of antidromic spikes elicited by stimulation of different forebrain sites. The latency of antidromic responses was shorter for sub-cortical stimulation sites compared to the cortical sites and latencies for more posterior cortices were longer in comparison to more anterior cortices. This is consistent with the LC projections, which pass through thalamus before entering anterior cortex and then traveling to the posterior cortex ^115^. (E) The mean spike rate of LC single units did not depend on their projection target, although there was a tendency for PFC-projecting units to spike at a higher rate than M1-projecting units (M1 v.s. ACC: T(9)=-2.18, p=0.063; M1 v.s. PL: T(20)=-1.07, p=0.296; M1 v.s. IL: T(18)=-2.282, p=0.035; M1 v.s. OFC: T(7)=-1.90, p=0.098). This result is in agreement with a recent study using LC slice recordings from neurons labeled with retrograde tracers injected in the OFC, PL, ACC, and M1 ^30^.

**Extended Data Figure 8.**
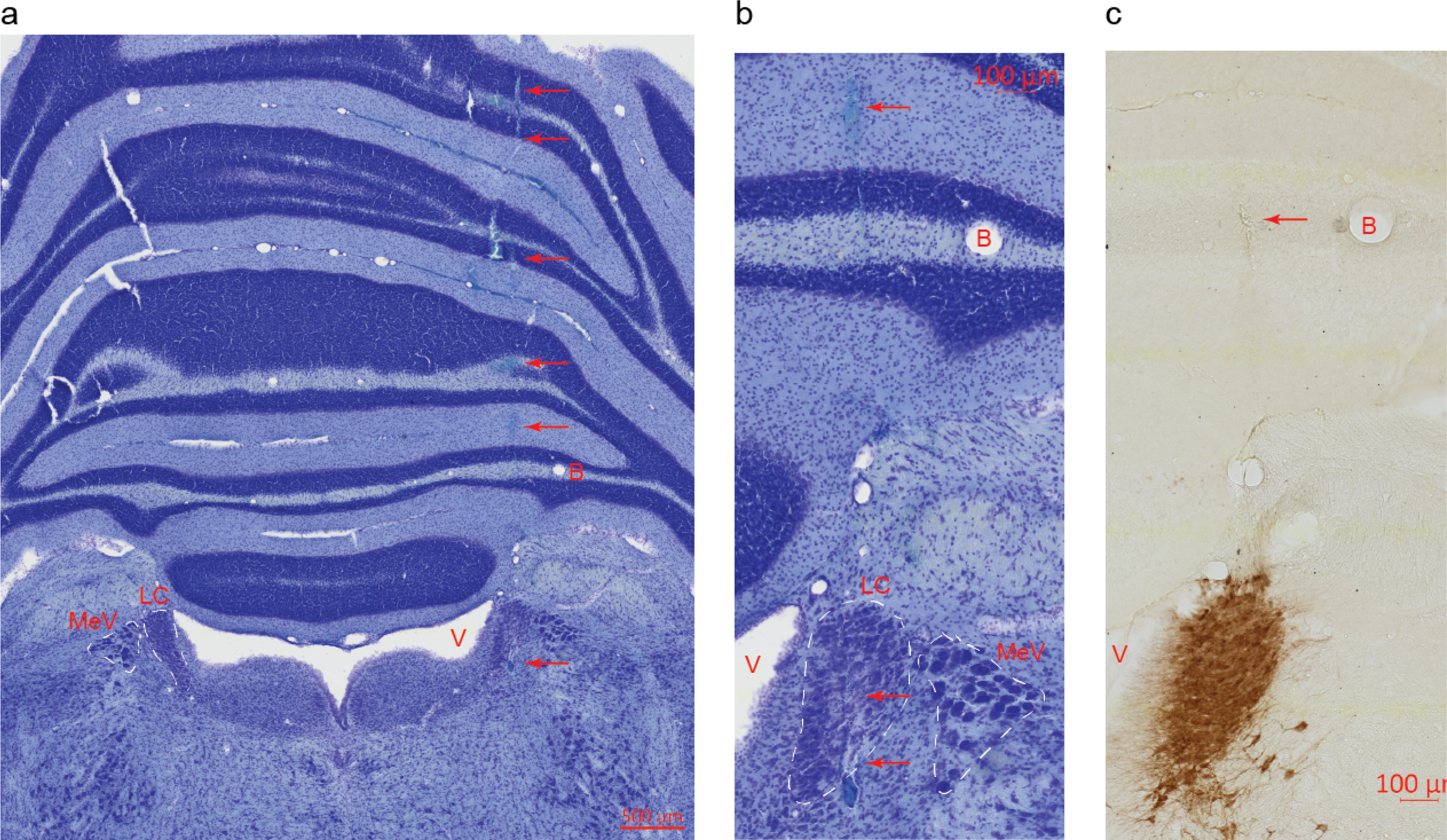
Histology illustrating an electrode track and the LC. A. Nissl stain in a coronal section (50 μm thickness) shows the electrode track made by the 15 μm thick probe within the coronal plane (arrows). Although accurate reconstruction of the electrode (15 μm thickness in the coronal plane) oriented in 50 μm thick coronal sections was difficult, the 15 degrees posterior angle of electrode insertion allowed visualization of the track dorsal to the LC traveling through the coronal plane. The dotted lines indicate the approximate extent of the LC and MeV (LC – Locus Coeruleus, MeV - Mesencephalic Nucleus of the Fifth Cranial Nerve, V – Fourth Ventricle). A blood vessel (B) is noticeable lateral to the electrode track and dorsal to the MeV. B. A close-up on the LC shown in A showing the LC and a track ventral to LC and medial to a blood vessel. C. The next section after the section in A and B that was stained with DAB against an antibody for the catecholamine-synthesis enzyme, Tyrosine Hydroxylase. Note the electrode track that is ventral to LC and medial to a blood vessel.

**Extended Data Table 1.** A list of the stereotactic coordinates for electrode placement. All coordinates are listed in millimeters.

| Region | Ref. | Anterior/Posterior | Lateral | Ventral from dura | Angle (deg) |
| --- | --- | --- | --- | --- | --- |
| LC | Lambda | -4.0 to -4.2 | 1.1 to 1.2 | 5.5 to 6.2 | 15 posterior |
| PL | Bregma | +3.0 | 0.6 | 3.0 | none |
| IL | Bregma | +3.0 | 0.6 | 4.4 | none |
| ACC | Bregma | +1.0 | 1.2 | 2.2 | 4 lateral |
| DLO | Bregma | +4.0 | 3.0 | 2.4 | none |
| M1 | Bregma | +3.0 | 3.0 | 1.6 | none |
| GI | Bregma | +1.0 | 5.0 | 4.0 | none |
| S1 | Bregma | -1.0 | 5.0 | 1.8 | none |
| S2 | Bregma | -2.0 | 4.5 | 4.15 | 15 lateral |
| BLA | Bregma | -1.88 to -2.12 | 4.6 to 4.9 | 6.8 | none |
| MDTh | Bregma | -3.0 | 1.0 | 5.0 | 2 lateral |
| Hb | Bregma | -3.0 | 1.0 | 4.5 | 2 lateral |
| VPMTh | Bregma | -3.0 | 3.0 | 5.6 | none |
| DLGN | Bregma | -4.0 | 3.5 | 4.0 | none |
| V2 | Bregma | -5.0 | 4.8 | 1.2 | none |
| V1 | Bregma | -7.0 | 3.0 | 1.4 | none |

